# Engineered endosymbionts that modulate primary macrophage function and attenuate tumor growth by shifting the tumor microenvironment

**DOI:** 10.1101/2024.07.18.604190

**Authors:** Cody S. Madsen, Ashley V. Makela, Chima V. Maduka, Emily M. Greeson, Anthony Tundo, Evran Ural, Satyajit H. Kulkarni, Ahmed Zarea, Matti Kiupel, Maryam Sayadi, Christopher H. Contag

**Affiliations:** Biosciences and Biotechnology Division, Lawrence Livermore National Laboratory, Livermore, CA, USA; Institute for Quantitative Health Science and Engineering, Michigan State University, East Lansing, MI, USA; Department of Biomedical Engineering, Michigan State University, East Lansing, MI, USA; Comparative Medicine & Integrative Biology, Michigan State University, East Lansing, MI, USA; Department of Microbiology Genetics and Immunology, Michigan State University, East Lansing, MI, USA; Department of Biochemistry and Molecular Biology, Michigan State University, East Lansing, MI, USA; Program in Cellular and Molecular Biology, Michigan State University, East Lansing, MI, USA; Department of Pathobiology and Diagnostic Investigation, Michigan State University, East Lansing, MI, USA

**Author notes:** BioFrontiers Institute, University of Colorado, Boulder, CO, USA. Institute for Quantitative Health Science and Engineering, and Department of Biomedical Engineering, Michigan State University, East Lansing, MI, USA. co-first authors.

**Keywords:** Engineered endosymbiont, transcription factors, bone marrow-derived macrophages, immune modulation, bacterial immunotherapy, tumor microenvironment, bacteriotherapy, immunometabolism

## Abstract

Modulating gene expression in macrophages can be used to improve tissue regeneration and to redirect tumor microenvironments (TME) toward positive therapeutic outcomes. We have developed *Bacillus subtilis* as an engineered endosymbiont (EES) capable of residing inside the eukaryotic host cell cytoplasm and controlling the fate of macrophages. Secretion of mammalian transcription factors (TFs) from *B. subtilis* that expresses listeriolysin O (LLO; allowing the EES to escape destruction by the macrophage) modulated expression of surface markers, cytokines and chemokines, indicating functional changes in a macrophage/monocyte cell line. The engineered *B. subtilis* LLO TF strains were evaluated in murine bone marrow-derived macrophages (BMDMs) by flow cytometry, chemokine/cytokine profiling, metabolic assays and RNA-Seq. Delivery of TFs by the EES shifted BMDM gene expression, production of cytokine and chemokines and metabolic patterns, indicating that the TF strains could guide primary macrophage function. Thereafter, the ability of the TF strains to alter the TME was characterized *in vivo*, in an orthotopic murine model of triple-negative breast cancer to assess therapeutic effects. The TF strains altered the TME by shifting immune cell composition and attenuating tumor growth. Additionally, multiple doses of the TF strains were well-tolerated by the mice. The use of *B. subtilis* LLO TF strains as EES showed promise as a unique cancer immunotherapy by directing immune function intracellularly. The uses of EES could be expanded to modulate other mammalian cells over a range of biomedical applications.

## Introduction

The concept of using endosymbionts and symbionts for modifying eukaryotic organisms to improve human health has been utilized in insects and nematodes^1–4^. *Wolbachia spp.* is a model endosymbiont that lives symbiotically within mosquitoes and naturally blocks the transmission of dengue and Zika virus by the mosquito species *Aedes aegypti*^5^ which has been used by researchers to stop transmission of these viruses in the environment^5,6^. Various symbionts have been engineered for applications ranging from improving honeybee immunity^7^ to enhancing nematode biocontrol of dangerous crop pathogens^8^. Furthermore, the benefits of symbiotic relationships between natural endosymbionts of invertebrates have been studied and have revealed clinically relevant compounds^9–15^. Previously, we developed and tested non-pathogenic engineered endosymbionts (EES) from a *B. subtilis* chassis that expresses listeriolysin O (LLO) and can persist in the cytoplasm of mammalian cells^16^. Variations of *B. subtilis* LLO expressing transcription factors (TFs) that altered macrophage function were delivered to a monocyte/macrophage cell line to study LLO-mediated entry into the host cell cytoplasm and modulation of macrophage pro- or anti-inflammatory responses^16^.

Bone marrow derived macrophages (BMDMs)^17,18^ represent primary, antigen-presenting cells that signal to other important immune cells and regulate immune response^19,20^ and may better represent the *in vivo* response than immortalized cell lines. Macrophages, including BMDMs, can function across a spectrum from being pro-inflammatory (M1) to anti-inflammatory (M2)^21–23^. These shifts in function can be distinguished by changes in gene expression, cell surface markers and expression of cytokines/chemokines^24–27^. Furthermore, shifts in cellular metabolism underlie macrophage polarization and immune function, such as after activation by lipopolysaccharide (LPS) or other pro-inflammatory stimuli ^28–38^. Changes in cellular metabolism can be revealed by measuring markers of transition from oxidative phosphorylation to glycolysis including change in oxygen consumption rate (OCR), extracellular acidification rate (ECAR), and adenosine triphosphate (ATP) production rate^39–43^. Metabolic shifts in macrophages occur as a component of the dynamic response to stimuli thus metabolic markers are indicative of macrophage responses. BMDMs have been used to study macrophage function in cancer^44–47^, chronic inflammation^48,49^, drug delivery^50,51^, pathogen response^52–54^ and tissue regeneration^51,55–57^.

Bacterial therapies based on extracellular bacteria have been developed to improve human health from improving the gut microbiome^58–61^ to treating cancer^62–65^. *Mycobacterium bovis* (Bacille Calmette-Guerin, BCG)^66,67^ was originally developed as a tuberculosis vaccine, but has now been approved for bladder cancer treatment, and other oncology applications are being tested^62–65,68–70^. Furthermore, several advancements have been made in improving extracellular bacteria treatment of cancer from tropism to therapeutic delivery^71–75^. *E. coli* Nissle 1917 (EcN) is a probiotic Gram-negative bacterium that has been part of the advancements in extracellular bacterial cancer immunotherapy^76–80^. EcN has been modified to deliver chemotherapeutic drugs and proteins while improving safety as a probiotic^75–79^. Intracellular bacteria have also been used in cancer immunotherapy. Gram-positive intracellular bacterium, *Listeria monocytogenes*, has been used to mobilize the immune system to alter the cancer microenvironment and *Salmonella typhimurium* has been used extensively to disrupt viability of cancer cells along with therapeutic molecule delivery^81–87^. Yet, these therapies are limited by dosed tolerance to live bacteria, especially Gram-negative bacteria, use of known pathogens as chassis organisms, and lack of characterized mechanisms on the target microenvironment^88–91^. *B. subtilis* LLO^92,93^, chassis of the EES, is derived from a non-pathogenic, generally recognized as safe (GRAS), Gram-positive, soil bacterium that respires as a facultative anaerobe and does not have a lipopolysaccharide-(LPS) mediated immune response which provides an alternative to some of the challenges described above^94,95^. The EES concept of using an intracellular approach to stimulate macrophages and mobilize the immune system to alter the cancer microenvironment parallels some efforts using *L. monocytogenes*^81,82^ for cancer treatment, but is built on a non-pathogenic bacterial platform that has been classically used for secretion of complex proteins^96^ and uniquely, it has been tied into mammalian cell regulatory pathways to redirect cell fates^16^.

In the therapies mentioned above, bacteria have been used to deliver a variety of therapeutic molecules including checkpoint inhibitors, nanobodies, or epitopes for vaccines, and have even been used to cause tumor cell lysis as a way of altering the tumor microenvironment (TME) or to activate immune cells^71–73,75,82^. TFs regulate genes that control cellular fates^97–100^ and thus TFs are being evaluated in clinical trials to treat cancer, and chronic wounds, as well as guide tissue regeneration and modulate immune responsesresponses^101^. Previously, we engineered *B. subtilis* LLO strains to express and deliver signal transducer and activator of transcription 1 (STAT-1) together with Krüppel-like factor 6 (KLF6) for polarization towards a pro-inflammatory phenotype, and Krüppel-like factor 4 (KLF4) together with GATA binding protein 3 (GATA-3) to polarize macrophages toward an anti-inflammatory phenotype^102–106^. Here we report that engineered *B. subtilis* LLO strains, expressing these TFs, altered patterns of BMDM gene expression, cytokine/chemokine expression and functional metabolism with patterns of modulation towards anti- or pro-inflammatory phenotypes within a complex response to the bacteria. Furthermore, in murine 4T1 orthotopic breast cancer the TME^107–109^ was altered by the engineered *B. subtilis* LLO strains with shifts in the immune cell composition leading to attenuation of tumor growth. Additionally, safety of this EES platform was observed as multiple doses could be injected without overt effects on the health of mice. By expressing TFs from the EES, macrophage function was modulated demonstrating that this approach is a promising therapeutic strategy and has applications in the study of cell biology (Supplementary Fig. 1-2).

## Results

### *B. subtilis* LLO escaped phagosome destruction, but were eliminated by autophagy

Fluorescent microscopy confirmed that *B. subtilis* LLO escaped phagosomal destruction in BMDMs (Fig. 1) with spatial overlap of bacteria (white) and LAMP-1^110^ positive structures (phagosomes, magenta)^111^. Phagosome escape was conditional on transcription induction by isopropyl β-D-1-thiogalactopyranoside (IPTG) of the *hlyA* gene encoding LLO which had also been observed for monocyte/macrophage cell lines^16,92^. Without IPTG addition (-IPTG), only very few intact *B. subtilis* rods were observed outside cells or as punctate regions within BMDMs, associated with LAMP-1 positive regions at 4 h (Fig. 1, orange arrow). By 12 h only remnants of bacteria were observed (Fig. 1, orange arrowhead). Conversely, when IPTG was added (+IPTG), *B. subtilis* rods were observed in several cells, not colocalized with LAMP-1 pockets, and some cells contained many bacteria at 4 h (Fig. 1, white arrow). Yet, after *B. subtilis* LLO escaped into the cytoplasm (+IPTG), BMDMs responded and removed most of the intracellular bacteria by 12 h (Fig. 1). Microtubule-associated protein light chain 3B (LC3) has been shown to coordinate autophagy response after phagosomal escape in macrophages infected with *L. monocytogenes* and other pathogens^112–115^. LC3B^112^ staining was used to indicate that this mechanism was activated at 12h by the BMDMs in response to intracellular *B. subtilis* LLO with some activation and destruction of *B. subtilis* LLO at 4 h (Fig. 1 white arrows). Accordingly, LLO expression when induced by IPTG, allowed *B. subtilis* LLO to access the cytoplasm of the BMDMs but resulted in *B. subtilis* LLO elimination within several hours. Nonetheless, live cell imaging validated that *B. subtilis* LLO remained viable within the cytoplasm as replication was observed in multiple cells, shown in a representative region of interest, between 3 and 4.5 h-post bacterial addition (Extended Fig. 1). Additionally, BMDMs were observed to actively pursue and share bacteria between cells to control bacterial proliferation and persistence (Extended Fig. 1).

**Fig. 1.**
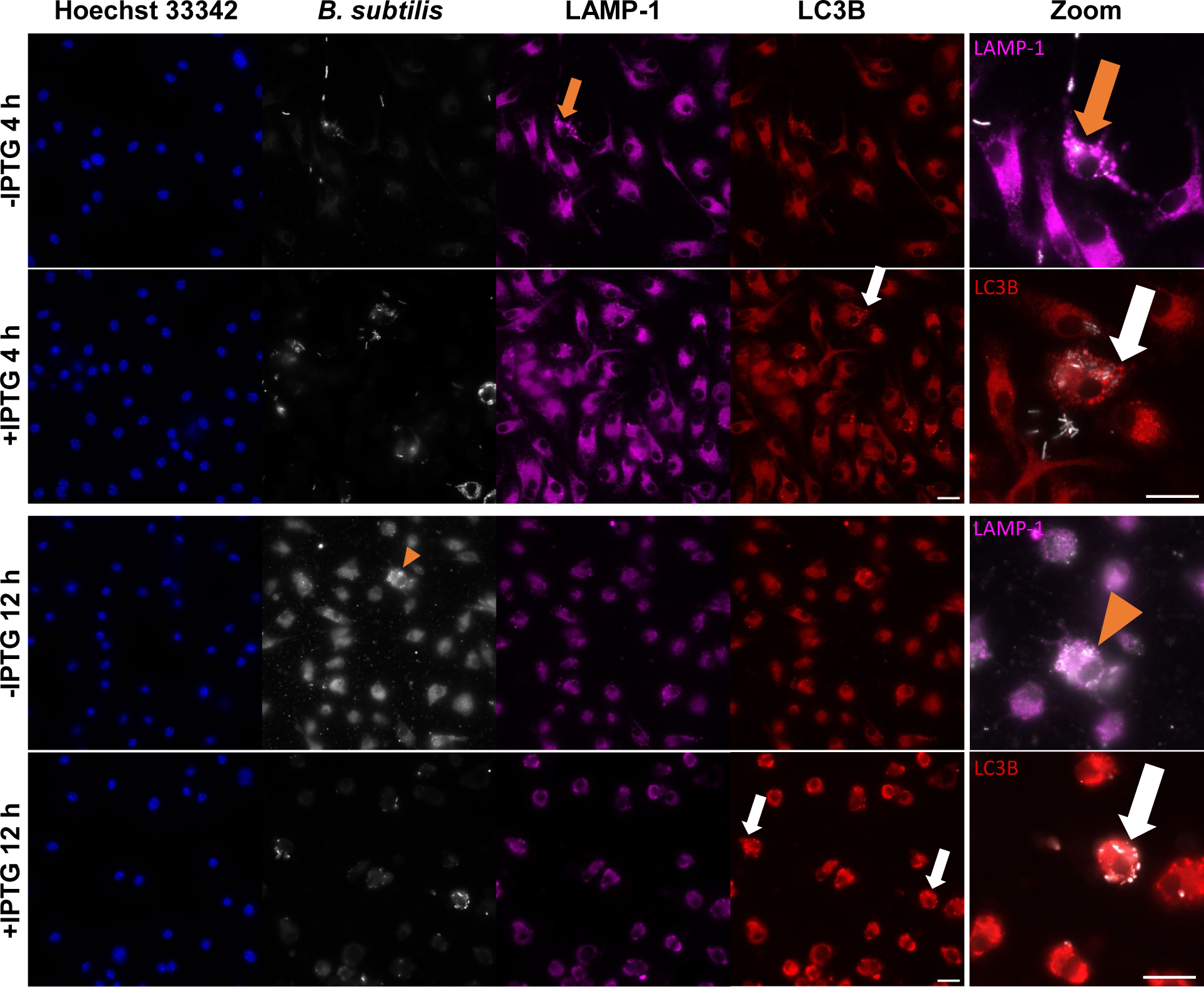
Fluorescent microscopy revealed *B. subtilis* LLO escape from BMDM phagosomes and destruction by autophagy.

BMDM autophagy after *B. subtilis* uptake was visualized using fluorescent microscopy. BMDM nuclei were stained by Hoechst 33342 (blue), and *B. subtilis* was stained by CellTracker Orange CMRA Dye (white). LAMP-1 (magenta) and LC3B (red) were used to stain subcellular structures. The LLO strain was delivered to BMDMs at a multiplicity of infection (MOI) of 50:1 and treated without (-IPTG) or with IPTG (+IPTG) with imaging at 4 and 12 h. Zoomed overlays of *B. subtilis* and LAMP-1 or LC3B are shown (right panel) for -IPTG and +IPTG conditions at 4 h (top) and 12 h (bottom). Without IPTG, there were few *B. subtilis* positive regions within cells and were located within LAMP-1 positive pockets in cells at 4 h (orange arrows). With IPTG (+IPTG), *B. subtilis* positive regions were present throughout several cells without strong LAMP-1 signal but, in some cells, positive LC3B pockets were associated with *B. subtilis* positive regions at 4 h (white arrows). By 12 h, only punctate *B. subtilis* positive regions were seen in the -IPTG condition, suggesting *B. subtilis* destruction, with few pockets of LAMP-1 and LC3B (orange arrowhead). However, addition of IPTG resulted *B. subtilis* positive regions at 12 h that were all associated with LC3B positive regions (white arrows). The z-depth was chosen for each overlayed image and each channel was adjusted to provide a representative image of each scenario. Scale bars = 20 µm.

### TF specific change of BMDM gene expression by engineered *B. subtilis* LLO strains

Macrophage gene expression patterns can reveal key behavioral changes. Gene expression is modulated by TFs as part of signaling cascades from various immune stimuli. Therefore, bulk RNA sequencing was performed to elucidate the BMDM gene expression changes in response to engineered *B. subtilis* LLO strains and controls. Data for all treatments was used to generate a 3D Principal Component Analysis (PCA) Emperor plot to visualize genome-wide gene expression changes for all treatments (Extended Fig. 2; https://github.com/madsen16/Engineered-endosymbionts). To assess the influence of TF on BMDM gene expression compared to the cytosolic presence of *B. subtilis* LLO, further PCA analysis was used to compare the TF strains to *B. subtilis* LLO. The TFs influenced genome-wide BMDM gene expression compared to the shifts caused by *B. subtilis* LLO (Fig. 2). Both *B. subtilis* LLO *Stat-1Klf6* (LLO-*SK*)^16^ and *B. subtilis* LLO *Klf4Gata-3* (LLO-*KG*)^16^ altered BMDM gene expression in patterns that were distinguishable compared to the LLO strain alone.

**Fig. 2.**
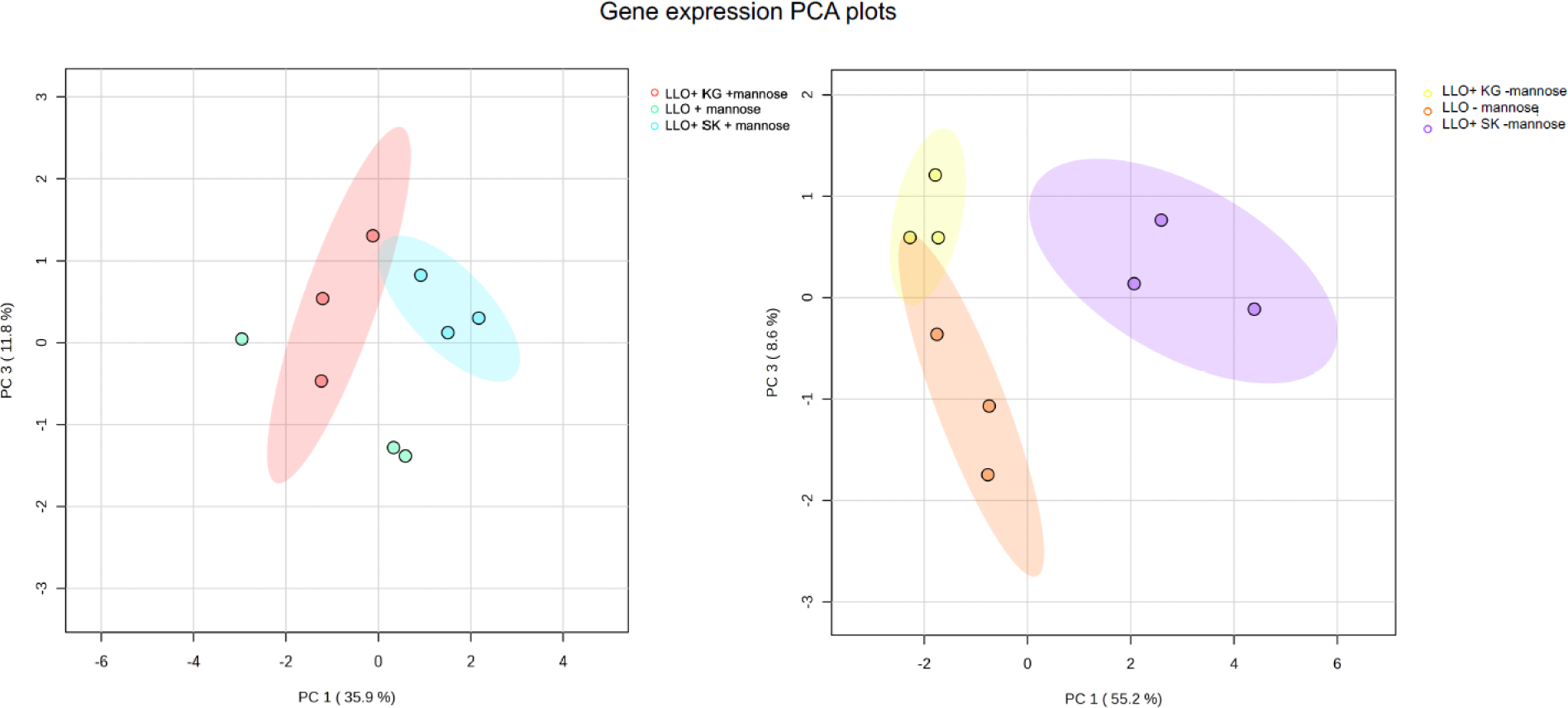
TFs directed BMDM gene expression.

PCA plots were used to visualize genome-wide gene expression differences between the following treatments: BMDMs cells treated with the LLO strain, with and without mannose (LLO -mannose, LLO +mannose), LLO-*SK* with and without mannose (LLO-*SK* -mannose, LLO-*SK* +mannose), and LLO-*KG* with and without mannose (LLO-*KG* -mannose, LLO-*KG* +mannose). The 24 h time points are shown. Shaded regions represent 95% confidence intervals.

Analysis of BMDM viability, marker expression and cytokine/chemokine production after exposure to engineered *B. subtilis* LLO strains

Flow cytometry was used to assess efficacy of *B. subtilis* LLO (LLO strain) internalization into BMDM *in vitro* when added at different MOI. *B. subtilis* LLO were stained with CellTracker Orange (CTO) CMRA Dye prior to co-incubation with BMDM, and added at MOI of 25:1, 50:1 or 100:1 (Extended Fig. 3). Cells which had CTO signal (%BMDM and alive) were identified as those containing *B. subtilis* LLO and the various MOI resulted in 16%, 33% and 29% positive cells, respectively. BMDM viability was also assessed by flow cytometry at an MOI of 50:1 at 4 and 12 h, either with or without LLO induction (±IPTG). After 4 h of incubation with the bacteria, IPTG induction of LLO caused a 5% loss of total cells with no significant difference in cell numbers at 12 h. When there was no LLO induction, no loss in BMDM viability was observed (Extended Fig. 3).

BMDM surface marker expression of CD11b+ F4/80+ was evaluated by flow cytometry to identify shifts in macrophage function caused by engineered *B. subtilis* LLO strains at 24 and 48 h. BMDMs were incubated with the LLO strain, LLO-*SK* or LLO-*KG*. For comparison, BMDMs were incubated in parallel with LPS and IFN-γ as a control for a pro-inflammatory phenotype, and IL-4 and IL-13 as an anti-inflammatory phenotype control. These were marked by expression differentiation markers CD86 and CD206^27,116,117^. For CD86, LPS and IFN-γ increased marker expression by 15- and 9-fold at 24 h and 48 h, respectively (Extended Fig. 4). There was a significant difference in CD86 expression among all treatment groups at both 24 h and 48 h time points (*p*=.016, Extended Fig. 4). BMDMs treated with all bacterial conditions, resulted in a trend of increased (4.92-fold mean) CD86 expression at 24 h when compared to untreated BMDM. At 48 h there was no significant increase in CD86 expression in any of the bacterial-treated groups. However, LLO-*KG* ± mannose treated groups had the greatest decrease in CD86 expression (average 2.7-fold less) versus LLO-*SK* ± mannose with the highest increase in CD86 expression (average 1.1-fold) when compared to untreated BMDM. For CD206, IL-4 and IL-13 increased marker expression by 6- and 2-fold at 24 h and 48 h, respectively (Extended Fig. 4). At 24 h there was no significant difference in CD206 expression in any bacteria treated groups compared to untreated BMDMs. At 48 h, all bacterial conditions caused a significant decrease in CD206 expression. There was no difference in CD206 expression between LLO strains, but there were increases in CD206 expression when mannose was added in the LLO (*p*=.101), LLO-*SK* (*p*=.011) and LLO-*KG* (*p*=.0008) groups.

As key mediators of immune cell signaling, cytokines and chemokines were profiled in BMDM cultures was to support translation to the *in vivo*^24,26^. BMDM cultures secreted significantly increased amounts of cytokines and chemokines in response to the bacterial treatments (several hundred or thousand-fold in some instances) and in many cases more than the positive controls, with LLO-*SK* and LLO-*KG* leading to different levels in the cultures (Fig. 3 Extended Fig. 5). LLO-*SK* and LLO-*KG* led to different levels of IL-10, macrophage inflammatory protein-1α (MIP-1α) and granulocyte colony stimulating factor (G-CSF) especially when TF expression was induced from the LLO strains by mannose; this was observed in comparison to each other, and when compared to the LLO strain at 24 h (Fig. 3a, e-f). The LLO-*KG* increased IL-6 levels in comparison to LLO-*SK* and the LLO strain at 24 h (Fig. 3b). Tumor necrosis factor alpha (TNF-α) increased with all bacterial treatments at 24 h, but at 48 h only the LLO-*SK* strain with mannose significantly increased TNF-α in comparison to the untreated BMDMs and comparable to that of the positive control (Fig. 3c-d). MIP-2 (CXCL2) and monocyte chemoattractant protein-1 (MCP-1/CCL2) showed specific responses to D-mannose with the different engineered strains still causing different patterns even with the complexity of addition of D-mannose at 24 h (Fig. 3g-h). D-mannose continued to impact the levels of these two proteins at 48 h (Extended 5e-f). While production of most cytokines and chemokines was triggered more by the bacteria than the positive controls, IL-12p40 and vascular endothelial growth factor (VEGF) did not show these trends. Production of both IL-12p40 and VEGF at 24 h and 48 h increased in response to the M1 positive control by either hundreds- or tens-fold, respectively (Extended Fig. 5m-p). The bacteria did not cause such large increases. Yet, at 24 h, IL-12p40 exhibited differential regulation by LLO-*SK* and LLO-*KG;* this was observed previously with LLO-*KG* increasing production by 5-fold in comparison to the untreated cells^16^. However, many of the cytokines and chemokines such as IL-10, IL-6 and MCP-1 are examples that did not show any differences in the bacterial treatments at 48 h (Extended Fig. 5a-f). Furthermore, some of the cytokines such as IL-15 and granulocyte-macrophage colony stimulating factor (GM-CSF) were not significantly produced in response to any treatments (Extended Fig. 5g-h, k-l). Finally, IL-1β was significantly produced only in response to the bacterial treatments at both 24 h and 48 h but no production difference was observed between the bacterial treatments (Extended Fig. 5i-j).

**Fig. 3.**
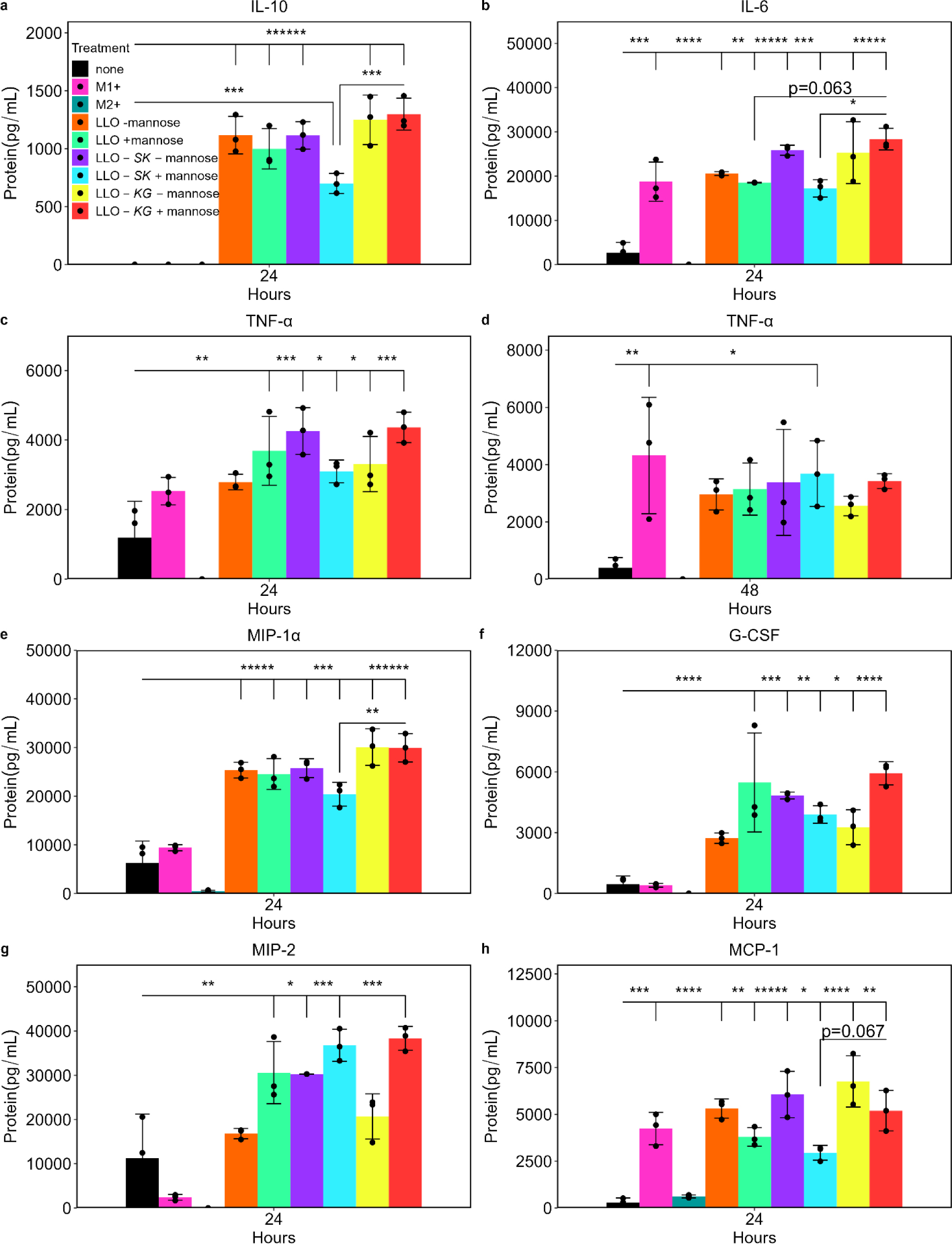
Engineered *B. subtilis* LLO strains altered BMDMs cytokine and chemokine production.

Cytokine and chemokine protein concentration was quantified in the media after BMDMs cells were untreated (none), treated with LPS and IFN-γ (M1+), IL-4 and IL-13 (M2+), LLO strain with and without mannose (LLO -mannose, LLO +mannose), LLO-*SK* with and without mannose (LLO-*SK* -mannose, LLO-*SK* +mannose) and LLO-*KG* with and without mannose (LLO-*KG* - mannose, LLO-*KG* +mannose) at 24 h and 48 h post-initial treatment. IPTG was added to all bacterial treatments. Data is mean ± SD from n=3 biological replicates; *p<0.05, **p<0.01, ***p<0.001, ****p<0.0001, *****p<0.00001, ******p<0.000001.

### Engineered *B. subtilis* LLO strains affect functional metabolism patterns in BMDMs maintained *in vitro*

Functional metabolism provides valuable insight into the activity and response of macrophages to various stimuli^118^. Additionally, D-mannose has been shown to impact macrophage metabolism and function^119^, therefore its effects were considered in this study. BMDM functional metabolism was characterized in response to the engineered *B. subtilis* LLO strains and mannose with LPS serving as a control, and these were compared to prior studies^32,33^. Overall, the D-mannose and engineered *B. subtilis* LLO strains modified BMDM functional metabolism in unique patterns at both 12 h and 24 h as indicated by OCR, ECAR and ATP production as indicators of shifts between glycolysis and oxidative phosphorylation (Fig. 4, Extended Fig. 6, Supplementary Table 1-2). The bacterial treatments increased basal OCR and ECAR similar to the LPS treatment at 12 h while results of D-mannose treatment were similar to no treatment (Fig. 4c-d). The LLO-*KG* strain, with and without mannose, increased basal OCR and ECAR more than any other condition at 12 h (Fig. 4c-d). Conversely, by 24 h, D-mannose overtook all other treatments as the dominating stimulus driving the change in metabolism even in the bacterial treatments (Extended Fig. 6c-d). The LLO-*KG* strain still caused trends of increased OCR and ECAR but only without the addition of D-mannose.

**Fig. 4.**
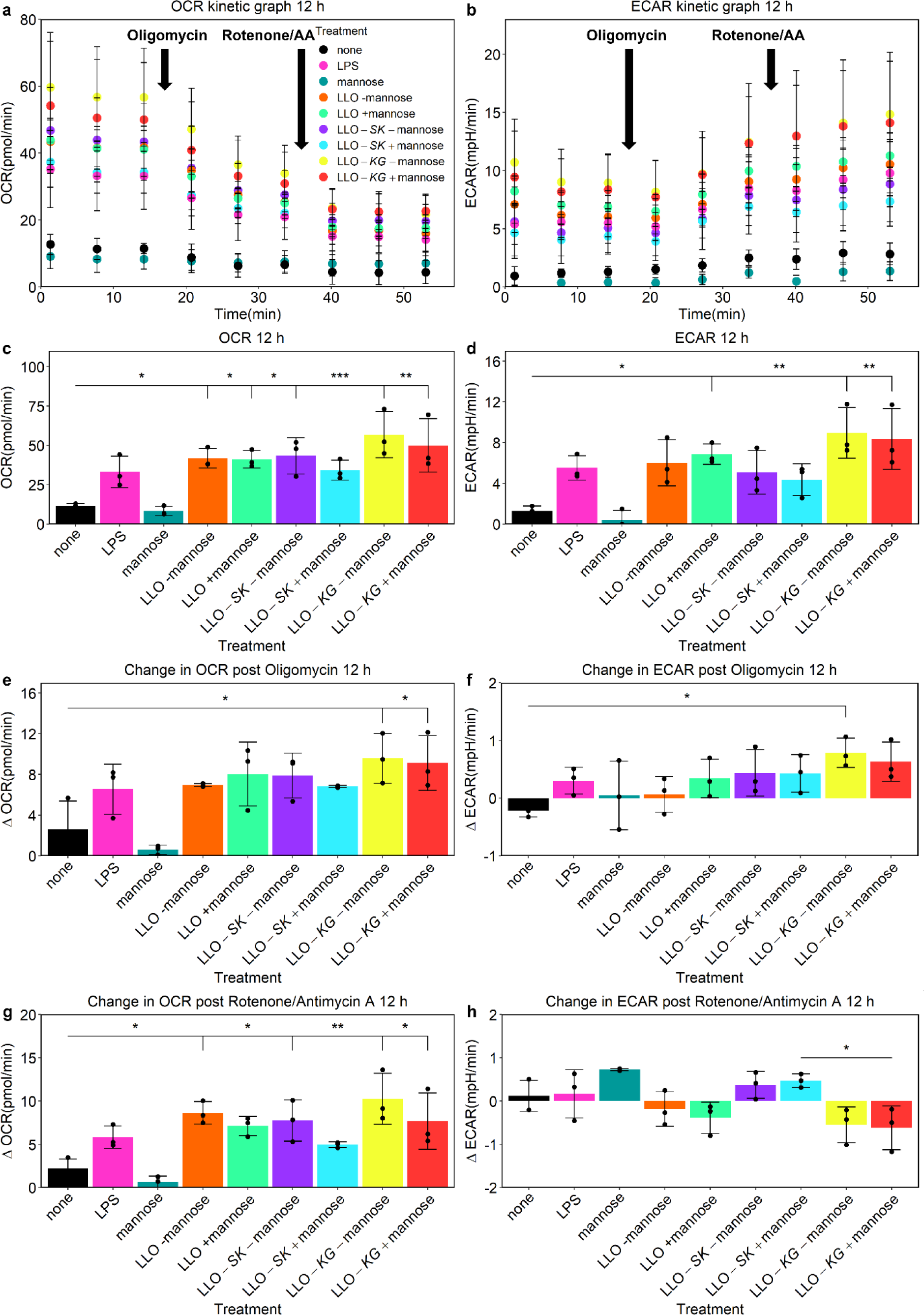
Shifts in functional metabolism patterns of BMDMs at 12 h after exposure to engineered *B. subtilis* LLO strains.

To further analyze responses to the bacteria alone and how the TFs may be impacting changes in metabolism, levels of OCR (ΔOCR) and ECAR (ΔECAR) were evaluated when the electron transport chain inhibitors oligomycin (complex V) and rotenone (complex I)/Antimycin A (AA; complex III) were added^120^. At 12 h, LLO-*KG* with, and without, D-mannose caused a significant change in OCR when Oligomycin was added, and Rotenone/AA treatment resulted in similar results as basal OCR (Fig. 4e, g). With changes in ECAR at 12 h, Rotenone/AA addition revealed different trends between the LLO-*SK* and LLO-*KG* strains after D-mannose treatment (Fig. 4h). At 24 h, D-mannose was the dominant stimulus when evaluating changes in OCR after inhibitors were sequentially added (Extended Fig. 6e, g). However, the changes in ECAR at 24 h after addition exhibited less dominance by D-mannose and even some differential trends between the engineered *B. subtilis* LLO strains when Oligomycin was added (Extended Fig. 6f, 6h).

Quantification of energy generation contribution from glycolysis or oxidative phosphorylation demonstrated that the bacterial treatments shifted the metabolism to glycolysis from oxidative phosphorylation at 12 h (Supplementary Table 1). The LLO-*KG* strain with mannose resulted in the most significant shift to glycolysis (*p*<0.007) comparable to that of LPS (*p*<0.005) while LLO-*SK* with mannose did not produce a significant shift. Additionally, total ATP production rates followed the same trends as the LLO-*KG* and LLO strain +mannose which increased ATP rates by 4-fold (*p*<0.01) and 3-fold (*p*<0.07), respectively, compared to the untreated cells at 12 h. D-mannose diverted all energy production to oxidative phosphorylation and led to a reduced ATP production rate at 12 h (Supplementary Table 1). Energetic contribution from either glycolysis or oxidative phosphorylation also paralleled the OCR and ECAR results at 24 h except in the LLO-*KG* with mannose treatment (Supplementary Table 2). The treatment of D-mannose at 24 h shifted energy generation towards glycolysis instead of oxidative phosphorylation which was inverse of trends at 12 h. The LLO-*KG* strain inhibited that shift when treated with D-mannose, which was similar to the other bacterial treatments without D-mannose. In contrast, treatment with the LLO strain and LLO-*SK* strain were significantly affected by the addition of D-mannose, p<0.04 and p<0.004, respectively (Supplementary Table 2). However, ATP production rates appeared to be largely driven by D-mannose in all bacterial treatments and were similar to that of the D-mannose treatment alone.

OCR and ECAR were measured before and after addition of the electron transport chain inhibitors, Oligomycin and Rotenone/antimycin A (AA), which were added at points indicated on kinetic plots (A, B). Basal OCR and ECAR quantification at the third measurement before addition of inhibitors were plotted (C, D). Further analysis was performed to quantify changes in OCR (ΔOCR) and ECAR (ΔECAR) after the inhibitors were added (E-H). BMDMs were untreated (none), treated with LPS, mannose, LLO strain with and without mannose (LLO -mannose, LLO +mannose), LLO-*SK* with and without mannose (LLO-*SK* -mannose, LLO-*SK* +mannose) and LLO-*KG* with and without mannose (LLO-*KG* -mannose, LLO-*KG* +mannose). IPTG was added to all bacterial treatments. Data is mean ± SD from n=3 biological replicates; *p<0.05, **p<0.01, ***p<0.001.

### Tumor growth rate decreased by engineered *B. subtilis* LLO strains without overt negative health effects on mice

Cancer bacteriotherapies have been shown to be most effective at reducing tumor growth and altering the tumor environment when injected intratumorally (IT)^72,73,88,121^. However, if the bacteria could be injected intravenously (IV) and were to naturally concentrate in the TME, this would facilitate clinical translation^122,123^. Therefore, we explored whether the engineered *B. subtilis* LLO strains would accumulate in the TME after IV injection and alter the TME. Bioluminescent, non-pathogenic *B. subtilis* LLO-*luxA-E*^121,122^ (LLO-*lux*) were shown to localize to 4T1 orthotopic tumors and persist in the tumor for a week after IV injection of 10^8^ bacteria while being cleared from healthy BALB/c mice in 24 h (Extended Fig. 7c, e & f). Furthermore, immunohistochemistry of excised tumor sections revealed Gram-positive structures (purple; black arrowheads) in the same spatial location as both phagocytes (CD11b+; brown; black arrow) and other CD11b-cells in the tumor (Extended Fig. 7g&h). Moving through the z-depth allowed for visualization of Gram-positive structures which were clumped together within cells. Tumor sections from mice, which did not receive a bacterial injection, did not have any detectable bacteria (Extended Fig. 7i). Accordingly, the effect on the progression of tumor growth was measured after the engineered *B. subtilis* LLO strains were injected. Animals received IV injections once a week for two weeks after which all mice were sacrificed and tumors were analyzed by TME immunophenotyping. LLO-*SK* +mannose caused the greatest reduction in normalized tumor growth (2.5-fold) relative to all other treatments, and in comparison, to the untreated animals, which was shown to be TF specific because all CFU counts were similar in all tumors (Extended Fig. 8). Furthermore, LLO-*SK* +mannose caused significant reduction in tumor growth as early as three days post-initial injection (Fig. 5). Also, LLO-*SK* +mannose significantly reduced tumor growth compared to LLO-*KG +*mannose, which reduced tumor growth by half in comparison to the untreated tumors (Fig. 5). In a separate group, LLO-*SK* was injected both IV (once a week) and IT (every third day between the weekly IV) to test for increased efficacy. LLO-*SK* IV+IT did not further reduce tumor growth in comparison to IV only (LLO-*SK;* Fig. 5). D-mannose alone also decreased tumor growth compared to the untreated tumors (Fig. 5, *p*<0.000001). When D-mannose was added to all bacterial treatments, only the TF strains showed a significant further reduction in tumor growth compared to the D-mannose treatment alone while the LLO-*lux* strain did not (Fig. 5).

**Fig. 5.**
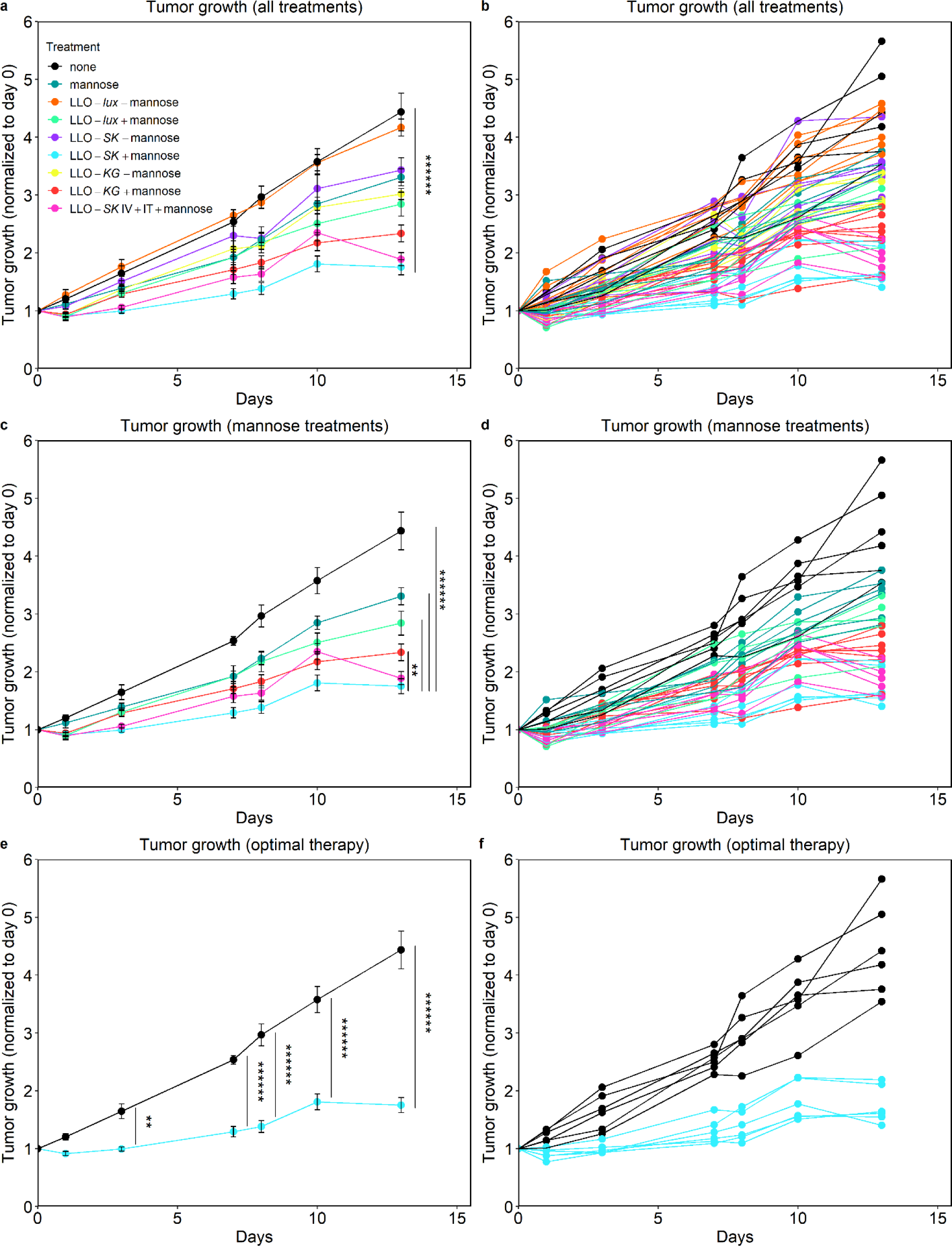
Bacteria-mediated reduction of tumor growth.

To investigate the health effects on the mice, mouse body mass was measured throughout the experiment and liver histopathology was used at the end of the experiment to determine any side effects in the liver. After intravenous injection of *B. subtilis* strains alone, there was no reduction in mouse body mass over the 13-day period. Only when D-mannose was added to drinking water did we observe a decrease in mouse body mass beginning at day 3. When compared to the untreated group, +mannose (*p*=0.013), LLO +mannose (*p*=0.001), LLO-*KG* +mannose (*p*=0.034) and LLO-*SK* IV+IT (*p*=0.011) groups had mouse body masses that were significantly lower at end point, but not the mice in the LLO-*SK* +mannose group. The group that lost the most weight by day 13 was the group which received mannose only (-9.2% loss compared to beginning of treatment). Liver histopathology did not identify any differences between untreated and bacteria treated groups. Within all groups there were multifocal perivascular and random mononuclear and neutrophilic cell infiltrates, and any other findings were normal within a physiological range of individual animals (Supplementary Table 3). Bacteria were not identified in the liver by light microscopy in any group following Giemsa staining. Additionally, after LLO-*lux* was injected into healthy mice, we observed zero CFUs in either the livers or the spleens of healthy, nontumor bearing, mice (Extended Fig. 8).

Tumor growth was measured in mm by calipers and calculated by the tumor volume equation then normalized to day of bacterial injection (day 0; tumors approximately 200 mm^3^ at day 0). Tumors were untreated (none), treated with mannose, LLO-*lux* strain injected IV with and without mannose (LLO-*lux* -mannose, LLO-*lux* +mannose), LLO-*SK* injected IV with and without mannose (LLO-*SK* -mannose, LLO-*SK* +mannose), LLO-*KG* injected IV with and without mannose (LLO-*KG* -mannose, LLO-*KG* +mannose) and LLO-*SK* injected IV and IT with mannose (LLO-*SK* IV+IT +mannose). IPTG was added to all bacterial treatments. Data is (a, c, e) mean ± SEM from n=6 mice; **p<0.01, ******p<0.000001.

### Tumor immunophenotyping and metabolism identified immune cell population and functional changes caused by engineered *B. subtilis* LLO strains

Profiling of the tumor immune cell populations to determine alterations to the TME showed that the engineered *B. subtilis* LLO strains can alter immune cell populations in tumors (Fig. 6, Extended Fig. 9). While there was no significant change in the total number of CD11b+ myeloid cells with bacterial treatment, when compared to the untreated group, bacterial treatments did decrease the number of monocytic myeloid-derived suppressor cells (M-MDSC; CD45^+^CD11b^+^Ly6C^high^Ly6G^-^) including treatments with LLO-*SK* multiple injections +mannose (*p*=0.0003) and LLO +mannose (*p*=0.032). Granulocytic myeloid-derived suppressor cells (G-MDSC; CD45^+^CD11b^+^Ly6G^+^Ly6G^low^) were decreased in the LLO-*SK* +mannose group when compared to +mannose only group (*p*=0.045) and LLO +mannose group (*p*=0.028). M2-like tumor associated macrophages (TAMs; CD45^+^CD11b^+^Ly6G^-^Ly6C^low^CD206^+^) decreased in the LLO-*SK* multiple injection +mannose group (*p*<.0001) and LLO +mannose group (*p*=0.041), which highlighted the ability to decrease these cell populations that are key players in tumor progression and contribute to a pro-tumoral phenotype. Further, the LLO-*SK* multiple treatment group significantly increased the M1/M2 TAM ratio compared to all other treatments, except LLO +mannose (*p*=0.086). Mature regulatory T cells (Tregs; CD25^+^FoxP3^+^) are another cell type known to suppress the immune response. Their presence was slightly decreased in the LLO-*SK* +mannose treated group (*p*=0.096) when compared to the untreated group. Furthermore, alternative FoxP3+CD25-populations, which are thought to have no suppressive function, were increased in the LLO-*SK* multiple injection group when compared to the untreated group (*p*=0.048)^124^. Tregs in the classical pre-activation state (FoxP3-CD25+) were decreased in the LLO-*SK* multiple injection group when compared to untreated (*p*=0.052) and the LLO +mannose group (*p*=0.047). It was seen that the introduction of LLO (without transcription factors or mannose) did not initiate a significant change in any immune cell population when compared to the untreated group. The introduction of mannose only influenced M MDSC (*p*<0.0001), when compared to untreated mice. There were no significant differences between LLO -mannose and LLO +mannose treatment groups in any of the analyzed immune cell populations. There were some changes in immune cell populations which were not expected; LLO-*SK* ±mannose and LLO-*KG* ±mannose all increased M2-like TAMs when compared to LLO +mannose. In addition, LLO-*SK* multiple injections decreased the dendritic cell (DC) population when compared to untreated mice (Fig. 6).

**Fig 6.**
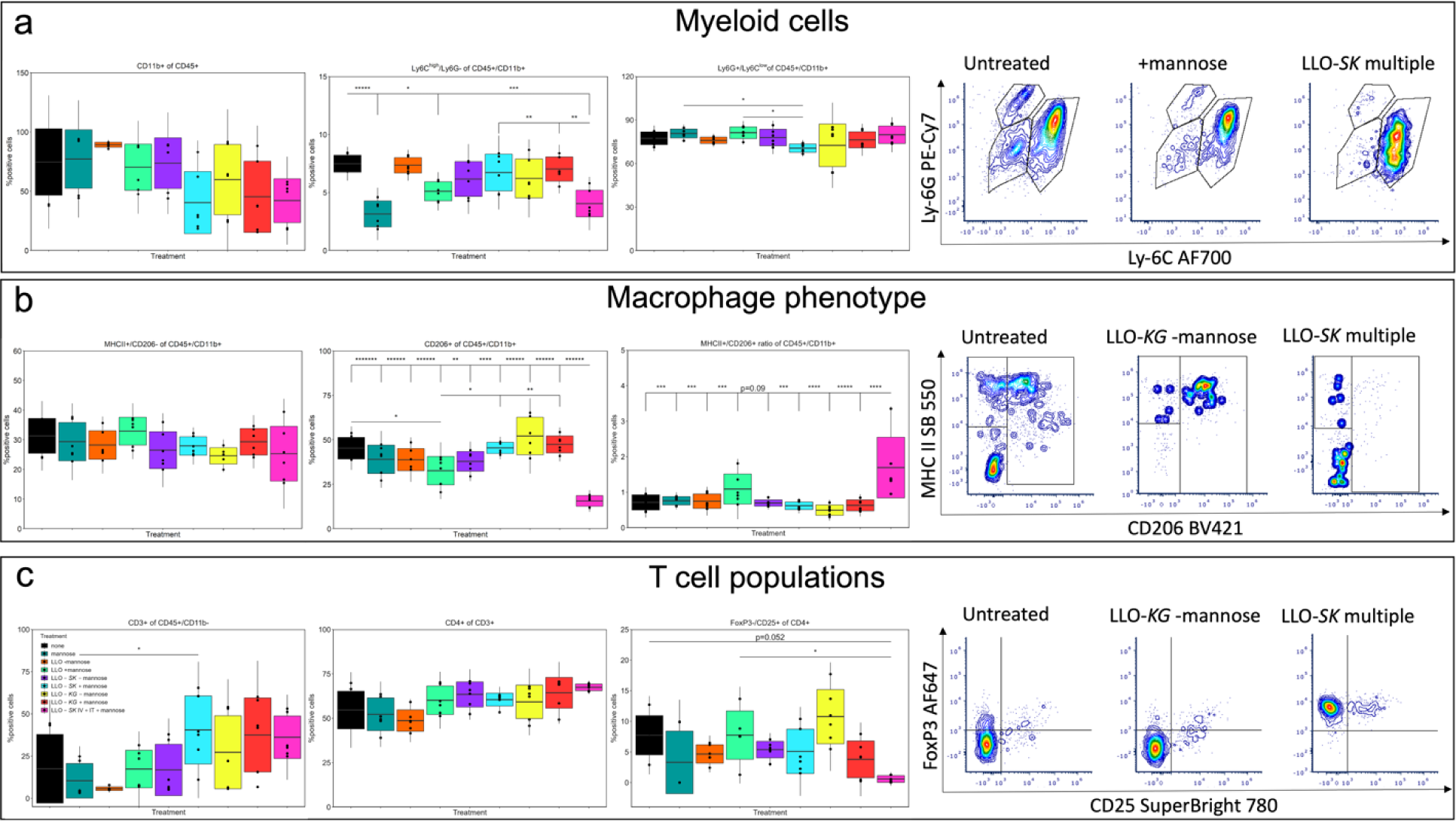
Tumor immunophenotyping revealed modifications in tumor immune cell populations.

Beyond profiling the immune cell populations, metabolism can be used to characterize changes to the TME as metabolism has been linked to cancer progression and to enhancing metastasis^36–38,120^. Combining this knowledge with the results that the engineered *B. subtilis* LLO strains could modulate functional metabolism, the TME metabolism was characterized^120^. We could associate mitochondrial bioenergetics within immune cell populations between individual mice in a treatment group and observed the same trend across groups. Both the LLO-*SK* +mannose and LLO-*KG* -mannose groups had the highest association between increased percent of CD3+ T-cells and increased basal OCR (R^2^=0.889, *p*=0.217 and R^2^=0.765, *p*=0.323, respectively; Supplementary Fig. 3). The group treated with mannose only, did not show any correlation (R^2^=0.002, *p*=0.975), suggesting that the presence of mannose in these groups did not have any effect on immunometabolism. A similar trend was observed in other measures such as max respiration as the LLO-*SK* +mannose treatment had the highest association between increased max respiration and increased CD3+ T-cells compared to other treatments such as LLO-*KG* -mannose and mannose (R^2^=0.898, *p*=0.207 and R^2^=0.82, *p*=0.279, R^2^=0.284, *p*=0.689, respectively).

Immune cell populations were analyzed using flow cytometry on cells from dissociated tumors in all treatment groups: untreated (none), treated with mannose, LLO-*lux* strain injected IV with and without mannose (LLO-*lux* -mannose, LLO-*lux* +mannose), LLO-*SK* injected IV with and without mannose (LLO-*SK* -mannose, LLO-*SK* +mannose), LLO-*KG* injected IV with and without mannose (LLO-*KG* -mannose, LLO-*KG* +mannose) and LLO-*SK* injected IV and IT with mannose (LLO-*SK* IV+IT +mannose). Populations analyzed were myeloid cells (CD11b^+^ of CD45^+^), monocytic myeloid derived suppressor cells (M MDSC; Ly6C^high^/Ly6G^-^ of CD11b^+^), granulocytic MDSC (G MDSC; Ly6G^+^/Ly6C^low^ of CD11b^+^), M1-like macrophages (MHCII^+^/CD206^-^ of CD11b^+^), M2-like macrophages (CD206^+^ of CD11b^+^), M2/M1-like population ratio, T cells (CD3^+^ of CD11b^+^), helper T cells (CD4^+^ of CD3^+^) and an alternative T regulatory cell subset (FoxP3^-^/CD25^+^ of CD4^+^).Data is mean ± SD for box and two times ± SD for whiskers from n=3 tumors in technical duplicates for each treatment; *p<0.05, **p<0.01, ***p<0.001, ****p<0.0001, *****p<0.00001, ******p<0.000001.

## Discussion

For the EES platform to be effective at altering macrophage function and to be used in cancer treatment, the EES must interact with the host cells and provide intracellular delivery of TFs. *B. subtilis* LLO gained access to the cytoplasm of BMDMs after IPTG induction of LLO expression (Fig. 1) and the cells were viable (Extended Fig. 1); this would allow for intracellular delivery of TFs. Even with destruction by LC3 related autophagy mechanisms^112–115^ bacteria could persist for several hours which seemed to be sufficient time for TF delivery, and perhaps after destruction could release TFs. Furthermore, the natural removal of the bacteria suggests safety of the EES platform when used as a therapy. Yet, if use of EES as a therapy requires more prolonged interaction between the host cell and engineered *B. subtilis* LLO strains, proteins such as phospholipases (plc) A and B from *L. monocytogenes* could be introduced into the *B. subtilis* LLO chassis to evade the autophagy^115^, but extended replication would need to be considered. After confirmed escape, replication and destruction by LC3 mechanisms of *B. subtilis* LLO, the viability of BMDMs remained high (∼95%), suggesting that macrophage function could be altered without loss of host cells (Extended Fig. 3). Yet, the incidence of uptake into BMDMs of approximately 35% resulted in an important consideration for other *in vitro* characterization and *in vivo* evaluation (Extended Fig. 3). The consideration included whether engineered *B. subtilis* LLO strains delivering TFs to 35% of the host cell population would result in functional change throughout the population and would that result in efficacy *in vivo*. Accordingly, the results seen throughout the study indicated that the cellular function, when measured in populations, was uniquely altered by the different bacterial strains.

Once the EES is in the cytoplasm, effective delivery of TFs, or other payloads designed to alter host cell gene expression, to the appropriate subcellular compartment is essential for functional change. The engineered *B. subtilis* LLO strains changed BMDM gene expression in patterns consistent with effective TF delivery, even when compared to the stimulus of *B. subtilis* LLO strain in the cytoplasm (Fig. 2). Overlap in genome-wide gene expression was observed more in the mannose conditions of the engineered *B. subtilis* LLO strains because of the effect of mannose on the BMDMs which has been demonstrated previously^119^ This observation is further supported by the conditions without mannose having further separation in these data plots, even without bacterial transcription of the TFs fully induced. Previously, the engineered *B. subtilis* LLO strains were shown to deliver TFs without transcription fully induced likely due to transcriptional leak in the mannose regulatory system^16,125^ and the partial effects of the TFs was still observed as indicated by alterations in gene expression patterns. Further in-depth analysis should be performed in future studies to understand what regulatory mechanisms and pathways are influenced by TF delivery. Additionally, more comparative analysis could be performed among all the treatment groups to assess new target pathways to be used in influencing macrophage behavior through new TF delivery or other molecules of interest (Extended Fig. 2).

Differential BMDM marker expression, cytokine/chemokine production and rates of metabolism provide a profile of functional changes in macrophage function, and an overview of the impact on the BMDMs by the bacteria and the TFs. The engineered *B. subtilis* LLO strains did not increase CD86 marker expression at 24 h but did show minimal change at 48 h in comparison to untreated BMDMs. CD206 was decreased by the strains but only at 48 h with the LLO-*KG* treatment causing the largest decrease of CD86 expression (Extended Fig. 4). However, these responses were greatly reduced in comparison to the monocyte/macrophage cell line response previously reported^16^. The lack of CD86 expression in response to the *B. subtilis* LLO from the BMDMs was unexpected because Toll-like receptor 2 (TLR2) triggers inflammatory responses^126,127^ but a similar lack of CD86 marker expression has been observed when bacterial TLR agonists were used to stimulate initially resting BMDMs which was also observed for CD206^128^. Unstimulated BMDMs appeared to produce a large response in cytokines but did not change marker expression unless pre-activation was performed before addition of the bacterial TLR agonists^128^. Further, the CD206 expression response was more complex because CD206 is the mannose binding receptor^129–131^ and can act as a receptor for bacterial surface carbohydrates^132^. Cytokines and chemokines play essential roles in signaling between various immune cell populations^26,52,133,134^ and these proteins have been shown to exhibit pleiotropic effects in disease progression and in cell classification^47,135–144^. Yet, unraveling this complexity provides valuable insights into outcomes *in vivo* and understanding therapeutic potential. The engineered *B. subtilis* LLO strains stimulated production of many tumor relevant cytokines and chemokines from the BMDMs with the TFs causing specific alterations presumably due to changes in gene regulation (Fig. 3, Extended Fig. 5). Accordingly, the LLO strain alone caused significant production of cytokines and chemokines (Fig. 3, Extended Fig. 5). LLO-*SK* and LLO-*KG* differentially regulated IL-10, MIP-1α, G-CSF and IL-12p40 at 24 h with IL-10, G-CSF and IL-12p40 occurring in a predicted pattern based on known regulation^16,104,106,145–149^. Further examples of predicted regulation include both LLO-*KG* upregulating IL-6 at 24 h^150^ and LLO-*SK* downregulating MCP-1^151,152^. D-mannose appeared to have a similar but opposite effect on MIP-2 as on MCP-1 and all treatments with D-mannose upregulated MIP-2 indicating that D-mannose is playing a role in macrophage response. Overall, the engineered *B. subtilis* LLO strains generated production of beneficial cytokines and chemokines for altering the TME including TNF-α and IL-12p40^148,149,153^ but the pleiotropic effects of cytokines and chemokines such as IL-6 and MCP-1 add ambiguity as both are implicated in promoting T cell invasion into the TME but can also promote tumor angiogenesis when expressed by cancer cells^151,154^. Yet, the LLO-*SK* strain did downregulate some cytokines and chemokines implicated in poor tumor outcomes such as MIP-1α, MIP-2, G-CSF and IL-10 which is beneficial^146,155–158^.

With the complexities seen in marker and cytokine/chemokine expression, functional metabolism was measured to provide another indicator of macrophage phenotype in response to the engineered *B. subtilis* LLO strains and relevance in tumors. In totality, the bacterial treatments increased glycolytic flux, similar to LPS treatment^28,30–35,159^. The LLO-*KG* strain promoted the most significant shift in metabolism with increased OCR, ECAR and ATP production (Fig. 4), likely due to the activity of KLF4 and GATA-3. Both TFs are involved in macrophage metabolism and work with STAT-6 to induce nuclear peroxisome receptor proliferator-activated receptor γ (PPARγ) causing mitochondrial biogenesis which could result in increased OCR and ATP production rate^104,118,160–162^. Additionally, OCR, an index of oxidative phosphorylation, can be simultaneously elevated with glycolytic flux during inflammation, being linked to production of cytokines and chemokines, such as IL-10, IL-6 and TNF-α^160,161^. These complex regulatory networks likely explain how the LLO-*KG* strain significantly increased oxidative phosphorylation and glycolysis at 12 h then was able to diminish a shift from oxidative phosphorylation to glycolysis caused by D-mannose at 24 h. Uniquely, OCR in the LLO-*KG* strain was consistently, significantly reduced by inhibition of complex V and complex I/ III of the ETC, suggesting that oxygen consumption is both directed at ATP production (complex V) and at superoxide formation (I/ III)^161^. Interestingly, a compensatory increase in ECAR was observed after complex I/ III inhibition, consistent with their interdependence^162^. Moreover, elevated ECAR with complex I/ III inhibition could indicate a role for STAT-1 and KLF6 in inflammation, as complex I generates nicotinamide adenine dinucleotide (NADH) when in reverse electron flow for lactate dehydrogenase to utilize in converting pyruvate to lactate which contributes to ECAR through H^+^ production^163,164^. D-mannose paralleled this result because D-mannose contributes to lactate production in macrophages^119^. The reduction in OCR, ECAR and ATP production with a significant shift towards glycolysis (except in LLO-*KG* condition) at 24 h in the D-mannose treatments could have resulted from the suppression of glucose utilization^119^ (Extended Fig. 6, Supplementary Table 2). TAMs have been shown to enhance tumor progression through metabolic changes^164–166^ and rely on glycolysis for energy production, so therapeutic measures have been proposed to increase PPARγ induction to cause increased phagocytic activity, which the LLO-*KG* caused in this study^164–166^. However, TAM metabolism is understudied and complex, so further studies are needed to understand the ideal target to adjust TAM metabolism^164–166^. Both the LLO-*SK* and LLO-*KG* strains caused functional changes that could be beneficial in TME altering applications with the LLO-*SK* strain expected to be most efficacious.

The limited examples of efficacy of bacteria immunotherapies when treating late-stage cancers have been related primarily to using pathogenic chassis organisms such as *S. typhimurium* due to direct impact on cancer cell viability^167,168^. However, when progressing to non-pathogenic chassis organisms and strategies involving activation of the immune system, late-stage cancers (e.g., 4T1 models) have proven challenging to treat^71,169,170^. The TF strains especially LLO-*SK* are an advance in the use of non-pathogenic chassis organisms that activate the immune system for translatable cancer treatment with efficacy and safety. Initially, the non-pathogenic LLO-*lux* strain was used to visualize location of bacterial accumulation temporally and was observed to be cleared from healthy mice within 24 h which was also observed with no CFU (Extended Fig. 8). However, the bacteria persisted within the tumors for a week post-injection IV and were found to be associated with phagocytes which established the potential for the engineered *B. subtilis* LLO strains to be utilized to alter the TME (Extended Fig. 7). The LLO-*SK* caused the greatest inhibition of tumor growth, which was observed at three days after initial injection and continued throughout the experiment (Fig. 5). LLO-*SK* had improved efficacy compared to LLO-*KG* which was expected based on the design of the TF pairs and the *in vitro* results with the BMDMs. LLO-*SK* was injected both IV and IT to reduce tumor progression potentially further as suggested in previous studies ^72,73,88,121^. However, we observed that the dual injections did not improve outcomes relative to single IV injections of LLO-*SK,* which may be explained by immunophenotyping (Fig. 5).

Tumor immunophenotyping by flow cytometry provides important insights into the composition of immune cell populations and can be used as a measure of efficacy in therapies which are designed to promote immune cell invasion into the TME^124,171,172^. This method has been used in bacterial cancer treatment to understand the benefits of using bacteria to activate immune cells towards treating cancer^71^. Analysis of the CD45+ hematopoietic cell composition in the 4T1 tumors revealed that the CD11b+ myeloid cell population was the predominant population (average 63.58%) among all CD45+ cells, as seen previously in the 4T1 tumor model^173^. CD11b+ cells can act in an immunosuppressive manner, inhibiting cancer immunity initiated by T and NK cells^174^. A reduction in the immunosuppressive CD11b+ and MDSC populations could indicate a therapeutic effect. Although we did not observe a clear change in CD11b+ populations among all the treatment groups, there were reductions in M-MDSC by the LLO-*SK* multiple treatment and LLO +mannose groups (Fig. 6). Mannose alone also decreased the M-MDSC population, highlighting how it can contribute to the alteration of the TME and tumor regression^175,176^. Another CD11b+ population within the TME is TAMs. Commonly, TAMs are of the pro-tumoral, immunosuppressive phenotype (M2), acting to help promote tumor growth, and metastases. The predominance of M2 macrophages is associated with a poor outcome while M1 macrophages are generally anti-tumoral ^21,22^. Although we did not observe an increase in the M1 population of TAMs, we could identify a decrease in the M2 population in the LLO-*SK* multiple injection and LLO +mannose groups (Fig. 6).

T cells are another important immune cell population which could provide therapeutic benefits during tumor treatment and the LLO*-SK* +mannose did increase the total population of T cells. The total population of CD3+ T cells can be composed of the therapeutic CD4+ or CD8+ populations^174–178^. On the other hand, the CD4+ population can harbor Tregs (classically defined as CD25+FoxP3+) and in a tumor setting these cells can suppress the immune response resulting in an inhibition of the ensuing antitumoral immune response^177–179^. Tregs are often identified using CD25 as an activation marker^180^ and more recently, FoxP3 which is required for development of Tregs and indicates a highly immunosuppressive population^181^. Our results suggest that mice which were treated with LLO-*SK* +mannose had a decreased population of mature Tregs (CD4+CD25+FoxP3+). Multiple treatments of LLO-*SK* decreased classical precursor Tregs (CD25+FoxP3-), which could have contributed to the limitation of tumor growth in these groups (Fig. 6, Extended Fig. 10)^182,183^.

Tumor metabolism analysis was performed to further understand the processes in the immune modulation goal of the EES platform. Even in the complex TME, we observed an indication towards activated T cell metabolism. Among conditions that had increased T cell populations (CD3+) including tumors within individual groups and across groups, an increase in OCR, ECAR, and ATP production was measured (Supplementary Fig. 3). Activated CD4+ and CD8+ T cells are known to increase both oxidative phosphorylation and glycolysis resulting in increased ATP production during a pro-inflammatory phenotype indicating various treatments especially LLO-*SK* +mannose were able to increase activated T cells in the TME^184^. Although the LLO-*SK* multiple injection treatment group had one of the best therapeutic effects, so did the group which was also injected with LLO-*SK* +mannose but only via two IV doses. Examining immunophenotyping, one may expect the group with multiple injections to have the most promising therapeutic effect. However, it is important to note that the multiple injections seemed to promote changes in macrophage populations while the single injection stimulated beneficial changes in T cells. Further studies could be performed to examine causes for the immune population differences between a single injection and multiple injection method.

After characterizing negative health effects on the mice, only mice which received mannose or bacteria +mannose had a significant reduction in weight except for LLO-*SK* (Extended Fig. 9). However, on average, mice that received the *B. subtilis* treatment had a weight reduction of 2.3% while the mannose treatment alone reduced weight by 8% indicating mannose in drinking water was the major contributor to weight reduction. Additionally, liver histopathology did not identify any differences between mice bearing 4T1 tumors with no treatment, mannose only treatment and the bacterial treatments (LLO, LLO-*SK*, LLO-*KG*) with mannose indicating safety of the *B. subtilis* treatments even at the primary location of initial bacterial accumulation and clearing. This observation is further supported by the clearing of *B. subtilis* from these organs in healthy mice (Extended Fig. 8).

Future directions could include various treatment regiments being tested to potentially improve efficacy, and this could be extended to other cancer models. For future advancement of the EES in cancer therapy and other applications, an extensive library of TF pairings, beyond these two pairs, should be developed along with improved genetic regulation elements to optimize macrophage response and improve stability. The design of bacterial operons expressing mammalian TFs could be improved by eliminating tandem repeats, while also refining control of expressed TF with the aim of maintaining EES stability especially in long-term *in vivo* experiments. Overall, the EES technology has utility for manipulating mammalian cell function in a targeted way that currently shows efficacy and safety as a cancer therapy and could be expanded to other biomedical applications that require alteration of the immune system.

## Acknowledgements

The authors would like to thank Dr. Daniel A. Portnoy for the *B. subtilis* LLO strain and Dr. Lee Kroos for *B. subtilis* plasmid pDR111. The authors would also like to thank Drs. Matthew Bernard and Daniel Vocelle of the Michigan State University Flow Cytometry Core for flow cytometry aid, cytokine/chemokine profiling analysis and tumor immunophenotyping design; the MSU RTSF Genomics Core for performing RNA-seq, MSU ICER for computational support in RNA-seq analysis. The authors would like to acknowledge the James and Kathleen Cornelius Endowment, the MSU College of Engineering Dissertation Completion Fellowship that supported C. S. Madsen and the MSU College of Natural Sciences Dissertation Continuation and Completion Fellowships that supported E. M. Greeson. Part of this work was performed under the auspices of the DOE by Lawrence Livermore National Laboratory under Contract DEAC52-07NA27344 (LLNL-JRNL-853911). Supplementary Fig. 1-2 were created using BioRender.com.

## Author contributions

Dr. Cody S. Madsen conceptualized the use of the EES as a bacterial cancer treatment, developed all the EES constructs as the platform technology, jointly developed and performed all experiments, jointly developed and analyzed all data to make figures and was one of the primary authors of the manuscript. Dr. Ashley V. Makela jointly developed and performed all experiments, jointly developed and analyzed all data to make figures and was one of the primary authors of the manuscript. Dr. Chima V. Maduka undertook the functional metabolism assays and significantly contributed to the writing and editing of this manuscript. Dr. Emily M. Greeson significantly contributed to the development of the EES platform technology, jointly designed and conducted CFU experiments, jointly developed and significantly contributed to the writing and editing of this manuscript. Anthony Tundo and Evran Ural both contributed to performing the *in vivo* studies and significantly contributed to the writing and editing of the manuscript. Dr. Matti Kiupel reviewed the pathology of the liver samples and edited the manuscript. Dr. Maryam Sayadi developed the RNA-seq analysis pipeline and significantly contributed to writing and editing of this manuscript. Dr. Christopher H. Contag conceptualized the use of EES for directing cellular therapies, supervised the studies, contributed to the experimental design, provided the resources, reviewed data and edited the manuscript.

## Methods

### *B. subtilis* LLO constructs

*B. subtilis* expressing IPTG-inducible LLO was provided by Dr. Daniel Portnoy. *B. subtilis* LLO *Stat-1Klf6* (LLO-*SK*) and *B. subtilis* LLO *Klf4Gata-3* (LLO-*KG*) were constructed and utilized in our previous work^16^. These same strains were utilized in this study and the *B. subtilis* LLO *luxA-E* (LLO-*luxA-E*) strain was constructed using the same homologous recombination plasmid (pDR111^185^, a gift from Dr. Lee Kroos) to insert the *luxA-E* operon into the *amyE* locus using a natural competence protocol^186^. The construct was selected by spectinomycin then confirmed by PCR amplification out of the genome and bioluminescence imaging. The *luxA-E* operon was amplified from a transposon plasmid used for bioluminescent imaging^187^ then inserted in place of the *lacI* gene in the pDR111 plasmid using inverse PCR then Gibson cloning for constitutive expression from the Phyper-spank promoter. Accordingly, this construct was inserted in the *B. subtilis* LLO *amyE* locus (LLO-*luxA-E* or LLO-*lux*).

### Growth conditions for *B. subtilis* LLO, LLO-*luxA-E* and TF strains

*B. subtilis* strains were grown under the same conditions for all experiments. Each *B. subtilis* LLO construct was grown in Luria-Bertani Miller broth (LB) with the appropriate antibiotic. *B. subtilis* LLO was grown in LB with chloramphenicol (10 µg/mL) while *B. subtilis* LLO-*SK*, *KG* and *luxA-E* were grown with spectinomycin (100 µg/mL). The overnight cultures were grown for 16 h at 37°C and 250 RPM. The *B. subtilis* LLO-*SK*, *KG* and *luxA-E* strains were added to macrophages *in vitro* or injected *in vivo* without antibiotics because constructs were integrated into the genome.

### *B. subtilis* LLO, LLO-*SK* and LLO-*KG* addition to BMDMs *in vitro*

The following conditions were utilized to induce *B. subtilis* LLO, LLO-*SK* and LLO-*KG* delivery, unless otherwise described. BMDMs were sourced from male and female C57BL/6J mice (Jackson Laboratories) of 3-4 months based on previous established methods^188^ and maintained at 37°C and 5% CO2 in DMEM (ThermoFisher, MA, USA), supplemented with 10% fetal bovine serum (FBS), 1% penicillin-streptomycin and 100 U/mL recombinant macrophage colony-stimulating factor (M-CSF). BMDMs were maintained in these conditions for 7 days before being seeded at 5×10^4^ into 96-well plates or 7×10^5^-1×10^6^ in 6-well plates and allowed to adhere overnight without penicillin-streptomycin. The engineered *B. subtilis* LLO were added at an optimized MOI of 50:1 for all experiments besides the uptake rate experiment (described below), along with IPTG (500 µM) to induce expression of LLO with or without delivery of TFs. The bacterial strains and BMDMs were then co-incubated at 37°C and 5% CO_2_ for 1 h. BMDMs were then washed three times with PBS and new medium was added containing gentamicin (5 µM) to eliminate any remaining extracellular bacteria. Co-incubation continued until BMDMs were evaluated by various analyses at various time points with the BMDMs eliminating intracellular engineered *B. subtilis* LLO within 11 additional hours (Supplementary Fig. 1). Further details of TF delivery in specific experiments are described below.

### Live cell imaging

*B. subtilis* LLO was added to BMDMs as described above, using a 96-well black glass-bottom plate (50,000 cells/well; Perkin Elmer, cat# 6005430). *B. subtilis* LLO was centrifuged (10,000 x g) for 2 min then resuspended in CellTracker Orange CMRA Dye (CTO, Invitrogen, C34564, 2 µM) in PBS then incubated at 37°C and 250 RPM for 25 min. Afterwards, *B. subtilis* LLO was centrifuged (10,000 x g) and washed three times before adding to BMDMs. Live cell imaging was performed on a Leica Dmi8 Thunder microscope equipped with a DFC9000 GTC sCMOS camera and LAS-X software (Leica, Wetzlar, Germany). BMDMs were maintained at 37°C and 5% CO_2_ in Fluorobrite medium during the imaging session. Fluorescent images of CTO were acquired using a TRITC filter set. Brightfield and fluorescent images were acquired consecutively, using a 63x oil objective every 1.5 h starting at 3 h post-bacterial addition until 9 h post-bacterial addition. Z-stacks were taken at all time points at 0.4 µm steps to confirm *B. subtilis* LLO presence within cytoplasm. Zoomed in images were created using Fiji (ImageJ; version 1.53t) software.

### Confirming *B. subtilis* LLO phagosomal escape and destruction by LC3 mechanisms

CTO was used to confirm location of *B. subtilis* LLO after being added to BMDMs in 96-well black glass-bottom plate (50,000 cells/well; Perkin Elmer, cat# 6005430) as described above. After fixation with 4% paraformaldehyde (PFA), cells were permeabilized using 0.3% Triton X-100 (ThermoFisher) followed by a blocking step containing 0.3% Triton X-100 and 5% normal goat serum (ThermoFisher, cat#31872,). Primary antibodies were incubated at 4°C overnight followed by secondary antibodies incubated at room temperature (RT) for 2 h. Phagosome formation or destruction was shown by incubating an anti-Lamp-1^110^ primary antibody (1:100, AbCam, MA, USA, cat#ab25245) followed by a goat anti-rat IgG Alexa Fluor 647 secondary antibody (1:5000, ThermoFisher, cat#A-21247). Autophagy mechanisms triggered by LC3 were elucidated by incubating an anti-LC3B primary antibody (1:1000, AbCam, MA, USA, cat#ab192890) followed by a goat anti-rabbit IgG Texas Red secondary antibody (1:2000, ThermoFisher, cat#T-2767). Nuclei were counterstained by incubating cells with Hoechst 33342 (1 µg/mL) for 10 minutes (min) at RT. Plates were imaged using a Leica Dmi8 Thunder microscope equipped with a DFC9000 GTC sCMOS camera and LAS-X software (Leica, Wetzlar, Germany). Brightfield and fluorescent images were acquired using a 63x oil objective with the fluorescent images acquired by the DAPI (Hoechst 33342), TRITC (CTO), Texas Red (LC3B) and Cy5 (LAMP-1) filter sets. Overlayed images were created using Fiji software.

### BMDM viability and uptake of *B. subtilis* LLO by flow cytometry

Flow cytometry was used to test for uptake of *B. subtilis* LLO and change in BMDM viability after bacterial delivery. For viability, conditions examined were multiple time points of interaction between *B. subtilis* LLO and host cells (4 and 12 h), a 50:1 MOI and with or without IPTG induction compared to untreated with biological triplicates (n=3) for each time and condition. At least 5,000 events were collected from the live gate for analysis. For uptake, conditions examined were multiple MOIs (25:1, 50:1, 100:1) compared to untreated after 4 h incubation with biological triplicate (n=3) for untreated and 50:1 MOI and one biological replicate (n=1) for 25:1 and 100:1 MOI. Cells were collected, washed once with 1X PBS and incubated with Zombie NIR viability dye (1:750, Biolegend, San Diego, CA, USA; cat#423105) in PBS for 20 min, at 4°C in the dark. Cells were washed twice followed by fixation using 4% PFA and resuspended in 100 µL flow buffer (0.5% bovine serum albumin (BSA), 1X PBS) for analysis using the Cytek Aurora Cytometer (Cytek Biosciences, CA, USA). All samples were assessed for percent live cells. BMDMs which were incubated with CTO bacteria were assessed for percent CTO positive cells (BMDMs containing bacteria), based on an FMO cutoff of 0.1% for CTO. Standard one-way ANOVA with Tukey post-hoc test was used to determine statistically different values.

### Engineered *B. subtilis* LLO strains TF delivery and BMDM protein production modulation

The LLO strain, LLO-*SK* and LLO-*KG* were internalized into BMDMs as described above (*B. subtilis* LLO, LLO-*SK* and LLO-*KG* addition to BMDMs *in vitro*), using a 6-well plate (Corning Costar #3516). D-mannose (1% w/v) was added to controls and to induce TF delivery after the initial 1 h coincubation between BMDMs and bacteria. TFs were delivered throughout the survival of the LLO-*SK* and LLO-*KG* strains and trafficked until experiments were ended for analysis (12, 24 or 48 h). For flow cytometry, Accutase (Sigma, cat#A6964) with scraping was used to detach BMDMs for analysis (described below). For Luminex cytokine/chemokine profiling (Millipore Sigma, MA, USA), the supernatant was removed at both 24 h and 48 h and then analysis was performed to quantify cytokines/chemokines produced (described below). BMDMs that were untreated were at resting state. BMDMs were polarized with IFN-γ and LPS (M1+) at 50 ng/mL and 100 ng/mL respectively or IL-4 and IL-13 (M2+) at 20 ng/mL each to be used as positive controls. All bacterial strains were treated with and without D-mannose and with IPTG as described above. All treatment conditions were performed in biological triplicates (n=3).

### *In vitro* flow cytometry

After addition of the engineered *B. subtilis* LLO strains and controls, followed by incubation for 24 h or 48 h, BMDMs were collected and stained in a 96-well round bottom plate. All staining steps were performed in 100 µL volume at 4°C in the dark. Samples were first incubated with Zombie NIR viability dye (1:750, Biolegend) for 20 min. Cells were washed once with flow buffer, followed by incubation with TruStain FcX™ PLUS (anti-mouse CD16/32) Antibody (Biolegend, cat#156603; 0.25 µg/sample) for 10 min. Alexa Fluor® 647 anti-mouse CD86 Antibody (0.125 µg/sample; Biolegend; cat#105020) and FITC anti-mouse CD206 (MMR) antibody (0.1 µg/sample, Biolegend; cat#141703) were then added and incubated for 20 min. Cells were washed twice with flow staining buffer and fixed with 4% PFA for 10 min and resuspended in a final volume of 100 µL for flow cytometry analysis using the Cytek Aurora spectral flow cytometer (Cytek). Five thousand events were collected from the live gate for analysis with single stained controls and unstained controls for all conditions used to assess fluorescent spread and for gating strategies. FMO gates had a cutoff percentage of 0.1%. CD86 and CD206 expression were analyzed from CD11b+/F4/80+ cells. At least 5,000 events were collected from the live gate for analysis. Flow cytometry data was analyzed with the software FCSExpress (DeNovo Software, CA, USA). Normality was tested using the Shapiro-Wild test and when failed, a Kruskal-Wallis test with Dunn’s multiple comparisons was performed. When normality was identified, a standard one-way ANOVA with Tukey’s multiple comparisons test was used. Statistical tests were used to determine statistically different MFI values amongst all groups within each time point. The data presented herein were obtained using instrumentation in the MSU Flow Cytometry Core Facility. The facility is funded in part through the financial support of Michigan State University’s Office of Research & Innovation, College of Osteopathic Medicine, and College of Human Medicine.

### Luminex cytokine/chemokine profiling assay

Cell culture supernatant was stored at -20°C until use (2 weeks). Supernatant was analyzed for CCL2 (MCP-1), CCL3 (MIP-1a), CXCL2/MIP-2, G-CSF, GM-CSF IL-1β, IL-6, IL-10, IL-12p40, IL-15, TNF-α and VEGFα cytokine expression. Cytokine and chemokine levels of cell supernatants were measured using a MCYTOMAG-70K Mouse Cytokine Magnetic Multiplex Assay (Millipore Sigma) using a Luminex 200 analyzer instrument (Luminex Corp, USA) according to the manufacturer’s instructions. Standard one-way ANOVA with Tukey post-hoc test was used to determine statistically different values amongst all treatment groups.

### *In vitro* Seahorse functional metabolism assays

Engineered *B. subtilis* LLO strains were added to BMDMs in a 96-well plate as described above (Engineered *B. subtilis* LLO strains TF delivery and BMDM protein production modulation). LPS (100 ng/mL) and D-mannose (1% w/v) served as controls in this experiment. Basal measurements of OCR and ECAR were obtained in real-time using the Seahorse XFe-96 Extracellular Flux Analyzer (Agilent Technologies) and was normalized to cell number^35,42,189^. Prior to running the assay, cell culture medium was replaced by the Seahorse XF DMEM medium (pH 7.4) supplemented with 25 mM D-glucose and 4 mM Glutamine. The Seahorse ATP rate and cell energy phenotype assays were run according to manufacturer’s instruction and all reagents for the Seahorse assays were sourced from Agilent Technologies. Wave software (Version 2.6.1) was used to process and export Seahorse data. Standard one-way ANOVA with Tukey post-hoc test was used to determine statistically different values amongst all treatment groups.

### RNA-sequencing

Engineered *B. subtilis* LLO strains were added to BMDMs in a 96-well plate as described above (Engineered *B. subtilis* LLO strains TF delivery and BMDM protein production modulation). The following treatments were used at concentrations described above in biological triplicate at both 12 h and 24 h (n=3): untreated, D-mannose, LPS, LPS and IFN-γ (M1+), IL-4 and IL-13 (M2+), LLO without IPTG (does not escape phagosomes), LLO strain with and without mannose (LLO - mannose, LLO +mannose), LLO-*SK* with and without mannose (LLO-*SK* -mannose, LLO-*SK* +mannose) and LLO-*KG* with and without mannose (LLO-*KG* -mannose, LLO-*KG* +mannose). These conditions and time points totaled 72 samples of mouse total RNA that was extracted by a Qiagen RNeasy kit (Qiagen, cat#74104) with RNase-free DNase Set (Qiagen, cat#79254) for NGS library preparation and sequencing. Libraries were prepared using the Illumina TruSeq Stranded mRNA Library Preparation Kit with IDT for Illumina TruSeq Unique Dual Index adapters following manufacturer’s recommendations. Completed libraries were quality controlled and quantified using a combination of Qubit dsDNA HS and Agilent 4200 TapeStation HS DNA1000 assays. The libraries were pooled in equimolar quantities for multiplexed sequencing. The library pool was loaded onto one lane of a NovaSeq S4 flow cell; sequencing was performed in a 2×150bp paired end format using a NovaSeq 6000 v1.5 300 cycle reagent cartridge. Base calling was done by Illumina Real Time Analysis (RTA) v3.4.4 and output of RTA was demultiplexed and converted to FastQ format with Illumina Bcl2fastq v2.20.0. All samples reached >30 million read counts. The quality of the reads was evaluated using FastQC (v. 0.11.7). All adaptors were trimmed using Trimmomatic (v. 0.39). The reads were aligned to GRCm39 genome using STAR (v. 2.6.0c). Reads were summarized using featureCounts and differential gene expression analysis was performed using Deseq2 (v.3.17).

### *In vivo* 4T1 tumor model and tumor growth measurements

Female BALB/c mice (6-8 weeks; Jackson Laboratories USA) were obtained and cared for in accordance with the standards of Michigan State University Institutional Animal Care and Use Committee. Mice were anesthetized with isoflurane administered at 2% in oxygen followed by an injection of 200,000 4T1 cells (a gift from Dr. Paula Foster, Western University, n=6 per group (n=7 in LLO-*KG* +mannose group); 98% viability, measured using the trypan blue exclusion assay) suspended in 50 µL PBS into the 4^th^ (inguinal) MFP, as previously reported^190^. Two weeks post-cancer cell implantation (pi) engineered *B. subtilis* LLO strains treatment was initiated. Mice were randomly divided into groups and separated based on strain injected and presence of D-mannose: 1) no treatment, 2) +mannose +IPTG, 3) LLO-*lux* intravenous (IV) -mannose +IPTG, 4) LLO-*lux* IV +mannose +IPTG, 5) LLO-*SK* IV -mannose +IPTG, 6) LLO-*SK* IV +mannose +IPTG, 7) LLO-*KG* IV -mannose +IPTG, 8) LLO-*KG* IV +mannose +IPTG, 9) LLO-*SK* IV/IT +mannose +IPTG. Bacterial treatments were administered under anaesthesia (as above). The engineered *B. subtilis* LLO strains were washed three times with PBS after overnight growth before resuspending at 1×10^8^ in 100 µL PBS then injected IV or resuspended and injected in 25 µL PBS for IT. After 24 h post-bacterial treatment, groups which were to be given D-mannose and IPTG received an IP injection of each (2 g/kg D-mannose, 50 mg/kg IPTG) and added to water (20% w/v D-mannose, 40 mM IPTG). Seven days after the first bacterial treatment, a second bacterial treatment was given, followed by D-mannose and IPTG 24 hours after. After the initial bacterial injection, animal well-being was documented every 2-3 days by observing water consumption, grooming and weight measurements. Tumors were measured using calipers beginning on the first day of the first treatment and continued every observation day of the second week until end point. Tumor volume was calculated using the equation^191^: tumor volume= 0.5(length×width^2^) from measurements taken by at least two individuals. A repeated measures two-way ANOVA with Tukey post-hoc test was used to determine any significance between treatments and time points. At endpoint, tumors were collected for flow cytometry immunophenotyping, histology, measurement of metabolism and colony forming units (below). Livers were also collected in formalin for histology to determine damage to tissue from repeated bacterial injections.

### Imaging of *B. subtilis* LLO-*lux* strains *in vivo*

The LLO-*lux* strain was imaged using the IVIS system (IVIS Spectrum, Perkin Elmer) using auto-exposure settings (time = 120-300 sec, binning = medium, f/stop = 1, emission filter = open). The LLO-*lux* strain was imaged immediately after injection, 1 h post-injection, 24 h post-injection, 72 h post-injection and at following regular time points that correlate with caliper measurements and animal well-being documentation. A final imaging time point was taken before euthanasia then after the tumors were removed and cut in half to elucidate location of bacteria throughout the tumor. Additionally, LLO-*lux* was injected into four mice (n=4) without tumors (-tumor) and imaged to track clearance of the bacteria using methods with tumor bearing mice mentioned above.

### Tumor immunophenotyping and metabolism characterization

Tumors were collected from each group (n=3) and halved for digestion into single cell suspension followed by immunophenotyping using flow cytometry analysis or metabolism characterization using the Seahorse Assay. Tumors were minced mechanically followed by digestion using a solution containing DMEM, Collagenase III (300 U/ml; Worthington Biochemical, cat#LS004182) and DNAse I (100 U/ml; Worthington Biochemical, cat#LS002139). Tumors were digested for 90 min at 37°C, 5% CO_2_ with agitation using a pipette every 30 min. The solution was passed through a 70 µm strainer followed by centrifugation and resuspension in ACK Lysing Buffer (ThermoFisher, cat#A1049201) for 1 min followed by addition of HBSS + 10% FBS. Cells were centrifuged, resuspended in PBS and counted using the Trypan Blue Assay. For immunophenotyping, 1×10^6^ cells (n=2) were collected per sample and transferred into a Nunc MicroWell 96-well polypropylene plate (Millipore Sigma, cat#P6866-1CS) for staining. All staining steps were performed in 100 µL volume at 4°C in the dark. Samples were first incubated with LIVE/DEAD Fixable Blue Dead Cell Stain (1:500, ThermoFisher, cat#L23105) for 30 min. Cells were washed once with flow buffer, followed by incubation with TruStain FcX™ PLUS (anti-mouse CD16/32) Antibody (Biolegend, cat#156603; 0.25 µg/sample) for 10 min. A mixture of the following antibodies was then added to the samples for 30 min: CD11b PE (0.125 ug/sample; Biolegend, cat#101207), CD8a BUV737 (0.25 µg/sample; ThermoFisher, cat#367-0081-80), CD25 SuperBright 780 (0.0625 µg/sample; ThermoFisher, cat#78-0251-82), CD3 APC/Fire 810 (0.25 µg/sample; Biolegend, cat#100267), Ly-6C PE/Cyanine7 (1:200; Biolegend, cat#128017), Ly-6G Alexa Fluor 700 (1:200; Biolegend, cat#127621), CD4 Brilliant Violet 510 (1:300; Biolegend, cat#100449), MHC-II (I-A/I-E) Spark Blue 550 (0.25 µg/sample; Biolegend, cat#107661), CD206 BV421 (1.25 µl/sample; Biolegend, cat#141717), NKp46 BV605 (1:100; Biolegend, cat#137619), CD11c BB700 (1:200; BD Bioscience, cat#566505), CD45 BUV395 (1:400; BD Bioscience, cat#564279) and CD19 BUV615 (0.125 ug/sample; BD Bioscience, cat#751213). Cells were washed followed by fixation and permeabilization for intracellular staining as per manufacturers protocol (eBioscience, cat#00-5523-00). FOXP3 AF647 (2.5 µl/sample; Biolegend, cat#320014) was then added to the samples for 30 min. Cells were washed twice with permeabilization buffer and resuspended in a final volume of 100 µL for flow cytometry analysis using the Cytek Aurora spectral flow cytometer (Cytek). Single stained controls and unstained controls for all conditions were used to assess fluorescent spread and FMO controls were used for gating strategies when needed. FMO gates had a cutoff percentage of 0.1%. At least 5,000 events were collected from the live gate for analysis. Flow cytometry data was analyzed with the software FCSExpress (DeNovo Software). For immunophenotyping, normality was tested using the Shapiro-Wild test and when failed, a Kruskal-Wallis test with Dunn’s multiple comparisons was performed. When normality was identified, a standard one-way ANOVA with Tukey’s multiple comparisons test was used. Statistical tests were used to compare all treatments.

For Seahorse analysis, 5×10^4^ cells from the halved and dissociated tumors in immunophenotyping above (n=3) were seeded into an appropriate 96-well plate in technical replicates of three wells (n=3). The Seahorse Mito Stress assay was run according to manufacturer’s instruction and all reagents for the Seahorse assays were sourced from Agilent Technologies. For the Seahorse analysis, a Brown-Forsythe and Welch ANOVA with Dunnett T3 post-hoc test was used to determine statistically different values amongst all treatment groups. For association analysis between immunophenotyping and metabolism, a linear regression model was used for statistical analysis.

### Colony forming units of *B. subtilis* LLO strains *in vivo*

After tumors were halved for immunophenotyping metabolism, two of the three alternate halves were used for CFU. Additionally, two whole tumors from the remaining three mice in the groups and one half of the final tumor were used. Whole tumors or halves were weighed then homogenized in 1 mL of PBS using a 1.5 mm Zirconium bead tube (Benchmark, cat#D1032-15) by bead beating in a Benchmark Beadbug 6 Microtube Homogenizer at max speed for 10 min. One hundred microliters of the homogenized mixture were then spread on LB plates with spectinomycin (100 µg/mL) for all groups and a 1:100 dilution was included for the LLO-*SK* IV+IT group. CFUs were counted and concentrations in tumors were calculated based on dilutions then normalized to tumor mass (g). The same process was repeated using whole livers and spleens from the three mice (n=3) used for immunophenotyping to calculate CFU in those organs including an additional three mice (n=3) from the -tumor group. Brown-Forsythe and Welch ANOVA with Dunnett T3 post-hoc test was used to determine any statistically different values amongst the treatment groups.

### Histological analysis

After the final time point, tumors and livers were collected and fixed in 4% PFA for 24 h. Tumors underwent cryopreservation through serial submersion in sucrose (10%, 20% and 30%). Tumors were frozen in optimal cutting temperature compound (Fisher HealthCare, USA) and sectioned using a cryostat (4 µm sections) and placed on a slide for staining. Sections were submerged in deionized water for 5 mins followed by blocking with 2% BSA in 1X PBS for 30 mins. Sections were then incubated with anti-CD11b polyclonal antibody (ThermoFisher, cat#PA5-79532, 1 ug/ml) for 2 h at room temperature. Sections were washed twice with 1X PBS followed by incubation with a secondary anti-rabbit IgG, HRP-linked antibody (Cell Signaling Technology, cat#7074S, 1:2000) for 30 mins. Sections were washed twice with 1X PBS followed by chromogenic detection using a DAB substrate kit (ThermoFisher, cat#34002) as per manufacturers suggestion. Sections were then stained to identify bacteria using a Gram Stain Kit (ThermoFisher, cat#R40080) and the following procedure: incubate with crystal violet (5 min), rinse with DI water, incubate with Gram’s iodine (2 min) and decolorize using 95% ethanol (30 sec). Sections were then dehydrated using 95% and 100% ethanol followed by clearing using xylene. Sections were imaged using a Nikon Eclipse Ci microscope equipped with a Nikon DS-Fi3 camera (Nikon, Tokyo, Japan) for color acquisition and NIS elements BR 5.21.02 software.

Formalin fixed samples of liver were sent to the MSU Veterinary Diagnostic Laboratory to be analyzed by a board-certified veterinary pathologist (Dr. Matti Kiupel). Briefly, samples were paraffin embedded after being processed in a Leica PELORIS II Premium Tissue Processing System using a standard 8 h xylene-free isopropanol schedule followed by sectioning (3 µm) and routine hematoxylin and eosin (H&E) and Giemsa staining.

### Statistical analysis and Reproducibility

Statistical analyses were performed using Prism software (9.4.0, GraphPad Inc., La Jolla, CA). Statistical tests are identified for each method. Data are expressed as mean +/- standard deviation unless stated otherwise; *p*<.05 was considered a significant finding. Plotting was performed using R version 4.0.4 with the following packages: ggplot2, dplyr, reshape2, ggsignif, plotrix and ggpubr.

## Data Availability statement

All raw data and engineered *B. subtilis* LLO constructs will be made available upon request by the corresponding author. Raw data used in the RNA-seq analysis can be accessed at Gene Expression Omnibus (GEO #GSE239519).

## Code Availability statement

All R scripts were written with a general format appropriate for the openly available, established packages mentioned above and can be made available on request. Access to 3D visualizations and code for RNA-seq analysis are available on GitHub (https://github.com/madsen16/Engineered-endosymbionts).

## Competing interests

The authors declare no competing interests.

## Notes

### Competing Interest Statement

The authors have declared no competing interest.

## References

1. Elston, K. M. et al. Engineering insects from the endosymbiont out. Trends Microbiol 30, 79–96 (2022).

2. Epis, S. et al. Chimeric symbionts expressing a Wolbachia protein stimulate mosquito immunity and inhibit filarial parasite development. Commun Biol 3, 1–10 (2020).

3. Wang, S. et al. Driving mosquito refractoriness to Plasmodium falciparum with engineered symbiotic bacteria. Science (1979) 357, 1399–1402 (2017).

4. Durvasula, R. V et al. Prevention of insect-borne disease: an approach using transgenic symbiotic bacteria. Proc Nat Acad Sci, USA 94, 3274–3278 (1997).

5. Jiggins, F. M. The spread of Wolbachia through mosquito populations. PLoS Biology Preprint at 10.1371/journal.pbio.2002780 (2017).

6. Flores, H. A. & O’Neill, S. L. Controlling vector-borne diseases by releasing modified mosquitoes. Nat Rev Microbiol 16, 508–518 (2018).

7. Leonard, S. P. et al. Engineered symbionts activate honey bee immunity and limit pathogens. Science (1979) 367, 573–576 (2020).

8. Machado, R. A. R. et al. Engineering bacterial symbionts of nematodes improves their biocontrol potential to counter the western corn rootworm. Nat Biotechnol 38, 600–608 (2020).

9. Xu, T.-T., Chen, J., Jiang, L.-Y. & Qiao, G.-X. Diversity of bacteria associated with Hormaphidinae aphids (Hemiptera: Aphididae). Insect Sci 28, 165–179 (2021).

10. Ip, J. C.-H. et al. Host-Endosymbiont Genome Integration in a Deep-Sea Chemosymbiotic Clam. Mol Biol Evol 38, 502—518 (2021).

11. Yang, K. et al. Wolbachia and Spiroplasma could influence bacterial communities of the spider mite Tetranychus truncatus. Exp Appl Acarol 83, 197–210 (2021).

12. Shi, X.-B. et al. Aphid endosymbiont facilitates virus transmission by modulating the volatile profile of host plants. BMC Plant Biol 21, 67 (2021).

13. Chao, L.-L., Castillo, C. T. & Shih, C.-M. Molecular detection and genetic identification of Wolbachia endosymbiont in Rhipicephalus sanguineus (Acari: Ixodidae) ticks of Taiwan. Exp Appl Acarol 83, 115–130 (2021).

14. Izraeli, Y. et al. Wolbachia influence on the fitness of Anagyrus vladimiri (Hymenoptera: Encyrtidae), a bio-control agent of mealybugs. Pest Manag Sci 77, 1023–1034 (2021).

15. Kennedy, J., Marchesi, J. R. & Dobson, A. D. W. Metagenomic approaches to exploit the biotechnological potential of the microbial consortia of marine sponges. Applied Microbiol Biotechnol Preprint at 10.1007/s00253-007-0875-2 (2007).

16. Madsen, C. S., Makela, A. V, Greeson, E. M., Hardy, J. W. & Contag, C. H. Engineered endosymbionts that alter mammalian cell surface marker, cytokine and chemokine expression. Commun Biol 5, 888 (2022).

17. Gonçalves, R. & Mosser, D. M. The Isolation and Characterization of Murine Macrophages. Curr Protoc Immunol 111, 14.1.1–14.1.16 (2015).

18. Zhang, X., Goncalves, R. & Mosser, D. M. The Isolation and Characterization of Murine Macrophages. Curr Protoc Immunol 83, 14.1.1–14.1.14 (2008).

19. Germic, N., Frangez, Z., Yousefi, S. & Simon, H.-U. Regulation of the innate immune system by autophagy: monocytes, macrophages, dendritic cells and antigen presentation. Cell Death Differ 26, 715–727 (2019).

20. Frucht, D. M. et al. IFN-γ production by antigen-presenting cells: mechanisms emerge. Trends Immunol 22, 556–560 (2001).

21. Mosser, D. M. & Edwards, J. P. Exploring the full spectrum of macrophage activation. Nat Revs Immunology Preprint at 10.1038/nri2448 (2008).

22. Moreira, A. P. & Hogaboam, C. M. Macrophages in allergic asthma: Fine-tuning their pro- and anti-inflammatory actions for disease resolution. J Interferon and Cytokine Res Preprint at 10.1089/jir.2011.0027 (2011).

23. Lavin, Y. et al. Tissue-Resident Macrophage Enhancer Landscapes Are Shaped by the Local Microenvironment. Cell 159, 1312–1326 (2014).

24. Mantovani, A. et al. The chemokine system in diverse forms of macrophage activation and polarization. Trends Immunol 25, 677–686 (2004).

25. Das, A. et al. High-Resolution Mapping and Dynamics of the Transcriptome, Transcription Factors, and Transcription Co-Factor Networks in Classically and Alternatively Activated Macrophages. Front Immunol 9, 22 (2018).

26. Turner, M. D., Nedjai, B., Hurst, T. & Pennington, D. J. Cytokines and chemokines: At the crossroads of cell signalling and inflammatory disease. Biochimica et Biophysica Acta (BBA) - Molecular Cell Research 1843, 2563–2582 (2014).

27. Martinez, F. O., Sica, A., Mantovani, A. & Locati, M. Macrophage activation and polarization. Frontiers in Bioscience Preprint at 10.2741/2692 (2008).

28. Liu, Y. et al. Metabolic reprogramming in macrophage responses. Biomark Res 9, 1–17 (2021).

29. Liu, D. et al. Comprehensive proteomics analysis reveals metabolic reprogramming of tumor-associated macrophages stimulated by the tumor microenvironment. J Proteome Res 16, 288–297 (2017).

30. Errea, A. et al. Lactate inhibits the pro-inflammatory response and metabolic reprogramming in murine macrophages in a GPR81-independent manner. PLoS One 11, e0163694 (2016).

31. Kelly, B. & O’neill, L. A. Metabolic reprogramming in macrophages and dendritic cells in innate immunity. Cell Res 25, 771–784 (2015).

32. Galván-Peña, S. & O’Neill, L. A. J. Metabolic reprograming in macrophage polarization. Front Immunol 5, 420 (2014).

33. Blagih, J. & Jones, R. G. Polarizing macrophages through reprogramming of glucose metabolism. Cell Metab 15, 793–795 (2012).

34. Lorenz, M. C., Bender, J. A. & Fink, G. R. Transcriptional response of Candida albicans upon internalization by macrophages. Eukaryot Cell 3, 1076–1087 (2004).

35. Ip, W. K. E., Hoshi, N., Shouval, D. S., Snapper, S. & Medzhitov, R. Anti-inflammatory effect of IL-10 mediated by metabolic reprogramming of macrophages. Science (1979) 356, 513–519 (2017).

36. Chen, Z., Lu, W., Garcia-Prieto, C. & Huang, P. The Warburg effect and its cancer therapeutic implications. J Bioenerg Biomembr 39, 267–274 (2007).

37. Escoll, P. & Buchrieser, C. Metabolic reprogramming of host cells upon bacterial infection: Why shift to a Warburg-like metabolism? FEBS J 285, 2146–2160 (2018).

38. Kim, J. & Dang, C. V. Cancer’s molecular sweet tooth and the Warburg effect. Cancer Res 66, 8927–8930 (2006).

39. Curi, R. et al. A past and present overview of macrophage metabolism and functional outcomes. Clin Sci 131, 1329–1342 (2017).

40. Lauterbach, M. A. et al. Toll-like receptor signaling rewires macrophage metabolism and promotes histone acetylation via ATP-citrate lyase. Immunity 51, 997–1011 (2019).

41. Pelletier, M., Billingham, L. K., Ramaswamy, M. & Siegel, R. M. Extracellular flux analysis to monitor glycolytic rates and mitochondrial oxygen consumption. in Meth Enzy vol. 542 125–149 (Elsevier, 2014).

42. Tannahill, G. M. et al. Succinate is an inflammatory signal that induces IL-1β through HIF-1α. Nature 496, 238–242 (2013).

43. Divakaruni, A. S., Paradyse, A., Ferrick, D. A., Murphy, A. N. & Jastroch, M. Chapter Sixteen - Analysis and Interpretation of Microplate-Based Oxygen Consumption and pH Data. in Mitochondrial Function (eds. Murphy, A. N. & Chan, D. C. B. T.-M. in E.) vol. 547 309–354 (Academic Press, 2014).

44. Cho, H. J. et al. Bone marrow-derived, alternatively activated macrophages enhance solid tumor growth and lung metastasis of mammary carcinoma cells in a Balb/C mouse orthotopic model. Breast Can Res 14, 1–12 (2012).

45. Weiser-Evans, M. C. M. et al. Depletion of cytosolic phospholipase A2 in bone marrow– derived macrophages protects against lung cancer progression and metastasis. Cancer Res 69, 1733–1738 (2009).

46. Redente, E. F. et al. Tumor progression stage and anatomical site regulate tumor-associated macrophage and bone marrow-derived monocyte polarization. Am J Pathol 176, 2972–2985 (2010).

47. Pathria, P., Louis, T. L. & Varner, J. A. Targeting Tumor-Associated Macrophages in Cancer. Trends Immunol 40, 310–327 (2019).

48. Davies, L. C. et al. Distinct bone marrow-derived and tissue-resident macrophage lineages proliferate at key stages during inflammation. Nat Commun 4, 1–10 (2013).

49. Redente, E. F. et al. Differential polarization of alveolar macrophages and bone marrow-derived monocytes following chemically and pathogen-induced chronic lung inflammation. J Leukoc Biol 88, 159–168 (2010).

50. Xia, Y. et al. Engineering Macrophages for Cancer Immunotherapy and Drug Delivery. Adv Materials 32, 2002054 (2020).

51. Spiller, K. L. & Koh, T. J. Macrophage-based therapeutic strategies in regenerative medicine. Adv Drug Deliv Rev 122, 74–83 (2017).

52. Kuhn, M. & Goebel, W. Induction of cytokines in phagocytic mammalian cells infected with virulent and avirulent Listeria strains. Infect Immun 62, 348–356 (1994).

53. Demuth, A., Goebel, W., Beuscher, H. U. & Kuhn, M. Differential regulation of cytokine and cytokine receptor mRNA expression upon infection of bone marrow-derived macrophages with Listeria monocytogenes. Infect Immun 64, 3475–3483 (1996).

54. Fortier, A., Faucher, S. P., Diallo, K. & Gros, P. Global cellular changes induced by Legionella pneumophila infection of bone marrow-derived macrophages. Immunobiol 216, 1274–1285 (2011).

55. Sun, D. et al. Bone marrow-derived cell regulation of skeletal muscle regeneration. The FASEB J 23, 382–395 (2009).

56. Gibon, E. et al. Aging affects bone marrow macrophage polarization: relevance to bone healing. Regen Eng Transl Med 2, 98–104 (2016).

57. Das, A. et al. Monocyte and macrophage plasticity in tissue repair and regeneration. Am J Pathol 185, 2596–2606 (2015).

58. Shen, T.-C. D. et al. Engineering the gut microbiota to treat hyperammonemia. J Clin Invest 125, 2841–2850 (2015).

59. Sheth, R. U., Cabral, V., Chen, S. P. & Wang, H. H. Manipulating Bacterial Communities by in situ Microbiome Engineering. Trend Gen 32, 189–200 (2016).

60. Puurunen, M. K., et al. Safety and pharmacodynamics of an engineered E. coli Nissle for the treatment of phenylketonuria: a first-in-human phase 1/2a study. Nat Metab 3, 1125–1132 (2021).

61. Charbonneau, M. R. et al. Development of a mechanistic model to predict synthetic biotic activity in healthy volunteers and patients with phenylketonuria. Commun Biol 4, 898 (2021).

62. Sedighi, M. et al. Therapeutic bacteria to combat cancer; current advances, challenges, and opportunities. Cancer Med 8, 3167–3181 (2019).

63. Sawant, S. S., Patil, S. M., Gupta, V. & Kunda, N. K. Microbes as Medicines: Harnessing the Power of Bacteria in Advancing Cancer Treatment. Int J Mol Sci 21, (2020).

64. Yaghoubi, A. et al. Bacteriotherapy in Breast Cancer. Int J Mol Sci 20, (2019).

65. Soleimanpour, S., Hasanian, S. M., Avan, A., Yaghoubi, A. & Khazaei, M. Bacteriotherapy in gastrointestinal cancer. Life Sci 254, 117754 (2020).

66. Zhang, Z. H., Yin, L., Zhang, L. L. & Song, J. Efficacy and safety of Bacillus Calmette-Guerin for bladder cancer: A protocol of systematic review. Medicine (Baltimore) 99, e21930 (2020).

67. Sfakianos, J. P. et al. Bacillus Calmette-Guerin (BCG): Its fight against pathogens and cancer. Urol Oncol 39, 121–129 (2021).

68. La Mantia, I. et al. The role of bacteriotherapy in the prevention of adenoidectomy. Eur Rev Med Pharmacol Sci 23, 44–47 (2019).

69. Andaloro, C., Santagati, M., Stefani, S. & La Mantia, I. Bacteriotherapy with Streptococcus salivarius 24SMB and Streptococcus oralis 89a oral spray for children with recurrent streptococcal pharyngotonsillitis: a randomized placebo-controlled clinical study. Eur Arch Otorhinolaryngol 276, 879–887 (2019).

70. Yaghoubi, A. et al. Bacteria as a double-action sword in cancer. Biochim Biophys Acta Rev Cancer 1874, 188388 (2020).

71. Chowdhury, S. et al. Programmable bacteria induce durable tumor regression and systemic antitumor immunity. Nat Med 25, 1057–1063 (2019).

72. Din, M. O. et al. Synchronized cycles of bacterial lysis for in vivo delivery. Nature 536, 81– 85 (2016).

73. Chien, T. et al. Enhancing the tropism of bacteria via genetically programmed biosensors. Nat Biomed Eng 6, 94–104 (2022).

74. Danino, T., Lo, J., Prindle, A., Hasty, J. & Bhatia, S. N. In vivo gene expression dynamics of tumor-targeted bacteria. ACS Synth Biol 1, 465–470 (2012).

75. Abedi, M. H. et al. Ultrasound-controllable engineered bacteria for cancer immunotherapy. Nat Commun 13, 1–11 (2022).

76. He, L. et al. Intestinal probiotics E. coli Nissle 1917 as a targeted vehicle for delivery of p53 and Tum-5 to solid tumors for cancer therapy. J Biol Eng 13, 1–13 (2019).

77. Yu, X., Lin, C., Yu, J., Qi, Q. & Wang, Q. Bioengineered Escherichia coli Nissle 1917 for tumour-targeting therapy. Microb Biotechnol 13, 629–636 (2020).

78. Danino, T. et al. Programmable probiotics for detection of cancer in urine. Sci Transl Med 7, 289ra84–289ra84 (2015).

79. Zhang, Y. et al. Escherichia coli Nissle 1917 targets and restrains mouse B16 melanoma and 4T1 breast tumors through expression of azurin protein. Appl Environ Microbiol 78, 7603–7610 (2012).

80. Luke, J. J. et al. Phase I Study of SYNB1891, an Engineered E. coli Nissle Strain Expressing STING Agonist, with and without Atezolizumab in Advanced Malignancies. Clin Can Res 29, 2435–2444 (2023).

81. Flickinger, J. C., Rodeck, U. & Snook, A. E. Listeria monocytogenes as a Vector for Cancer Immunotherapy: Current Understanding and Progress. Vaccines vol. 6 Preprint at 10.3390/vaccines6030048 (2018).

82. Morrow, Z. T., Powers, Z. M. & Sauer, J.-D. Listeria monocytogenes Cancer Vaccines: Bridging Innate and Adaptive Immunity. Curr Clin Microbiol Rep 6, 213–224 (2019).

83. Hayashi, K. et al. Cancer metastasis directly eradicated by targeted therapy with a modified Salmonella typhimurium. J Cell Biochem 106, 992–998 (2009).

84. Nagakura, C. et al. Efficacy of a genetically-modified Salmonella typhimurium in an orthotopic human pancreatic cancer in nude mice. Anticancer Res 29, 1873–1878 (2009).

85. Liang, K. et al. Genetically engineered Salmonella Typhimurium: Recent advances in cancer therapy. Cancer Lett 448, 168–181 (2019).

86. Nguyen, V. H. et al. Genetically engineered Salmonella typhimurium as an imageable therapeutic probe for cancer. Cancer Res 70, 18–23 (2010).

87. Guo, Y. et al. Targeted cancer immunotherapy with genetically engineered oncolytic Salmonella typhimurium. Cancer Lett 469, 102–110 (2020).

88. Harimoto, T. et al. A programmable encapsulation system improves delivery of therapeutic bacteria in mice. Nat Biotechnol (2022) doi:10.1038/s41587-022-01244-y.

89. Toso, J. F. et al. Phase I study of the intravenous administration of attenuated Salmonella typhimurium to patients with metastatic melanoma. J Clin Oncol 20, 142–152 (2002).

90. Yelin, I. et al. Genomic and epidemiological evidence of bacterial transmission from probiotic capsule to blood in ICU patients. Nat Med 25, 1728–1732 (2019).

91. Low, K. B. et al. Lipid A mutant Salmonella with suppressed virulence and TNFα induction retain tumor-targeting in vivo. Nat Biotechnol 17, 37–41 (1999).

92. Bielecki, J., Youngman, P., Connelly, P. & Portnoy, D. A. Bacillus subtilis expressing a haemolysin gene from Listeria monocytogenes can grow in mammalian cells. Nature 345, 175–176 (1990).

93. Zuber, P. & Losick, R. Role of AbrB in Spo0A- and Spo0B-dependent utilization of a sporulation promoter in Bacillus subtilis. J Bacteriol (1987) doi:10.1128/jb.169.5.2223-2230.1987.

94. Travassos, L. H. et al. Toll-like receptor 2-dependent bacterial sensing does not occur via peptidoglycan recognition. EMBO Rep 5, 1000–1006 (2004).

95. Elshaghabee, F. M. F., Rokana, N., Gulhane, R. D., Sharma, C. & Panwar, H. Bacillus As Potential Probiotics: Status, Concerns, and Future Perspectives. Front Microbiol 8, 1490 (2017).

96. Kolkman, M. A. B. et al. The twin-arginine signal peptide of Bacillus subtilis YwbN can direct either Tat- or Sec-dependent secretion of different cargo proteins: Secretion of active subtilisin via the B. subtilis Tat pathway. Appl Environ Microbiol 74, 7507–13 (2008).

97. Tugal, D., Liao, X. & Jain, M. K. Transcriptional control of macrophage polarization. Arterioscler Thromb Vasc Biol (2013) doi:10.1161/ATVBAHA.113.301453.

98. Mitchell, P. J. & Tjian, R. Transcriptional regulation in mammalian cells by sequence-specific DNA binding proteins. Science (1979) 245, 371–378 (1989).

99. Fleetwood, A. J., Lawrence, T., Hamilton, J. A. & Cook, A. D. Granulocyte-macrophage colony-stimulating factor (CSF) and macrophage CSF-dependent macrophage phenotypes display differences in cytokine profiles and transcription factor activities: implications for CSF blockade in inflammation. J immunol 178, 5245–5252 (2007).

100. Heinz, S. et al. Simple combinations of lineage-determining transcription factors prime cis-regulatory elements required for macrophage and B cell identities. Mol Cell 38, 576–589 (2010).

101. Rilo-Alvarez, H., Ledo, A. M., Vidal, A. & Garcia-Fuentes, M. Delivery of transcription factors as modulators of cell differentiation. Drug Deliv Transl Res 11, 426–444 (2021).

102. Li, H., Jiang, T., Li, M. Q., Zheng, X. L. & Zhao, G. J. Transcriptional regulation of macrophages polarization by microRNAs. Front Immunol Preprint at 10.3389/fimmu.2018.01175 (2018).

103. Date, D. et al. Kruppel-like transcription factor 6 regulates inflammatory macrophage polarization. J Biol Chem (2014) doi:10.1074/jbc.M113.526749.

104. Liao, X. et al. Krüppel-like factor 4 regulates macrophage polarization. J Clin Invest (2011) doi:10.1172/JCI45444.

105. Yang, M. et al. Deficiency of GATA3-Positive Macrophages Improves Cardiac Function Following Myocardial Infarction or Pressure Overload Hypertrophy. J Am Coll Cardiol (2018) doi:10.1016/j.jacc.2018.05.061.

106. VanDeusen, J. B. et al. STAT-1-mediated repression of monocyte interleukin-10 gene expression in vivo. Eur J Immunol 36, 623–630 (2006).

107. Tao, K., Fang, M., Alroy, J. & Sahagian, G. G. Imagable 4T1 model for the study of late stage breast cancer. BMC Cancer 8, 228 (2008).

108. Zhang, H. et al. Oncolytic adenoviruses synergistically enhance anti-PD-L1 and anti-CTLA-4 immunotherapy by modulating the tumour microenvironment in a 4T1 orthotopic mouse model. Cancer Gene Ther 29, 456–465 (2022).

109. Perez, G. I. et al. In Vitro and In Vivo Analysis of Extracellular Vesicle-Mediated Metastasis Using a Bright, Red-Shifted Bioluminescent Reporter Protein. Adv Genetics 3, 2100055 (2022).

110. Kortebi, M. et al. Listeria monocytogenes switches from dissemination to persistence by adopting a vacuolar lifestyle in epithelial cells. PLoS Pathog 13, e1006734–e1006734 (2017).

111. Huynh, K. K. et al. LAMP proteins are required for fusion of lysosomes with phagosomes. EMBO J 26, 313–324 (2007).

112. Deretic, V., Saitoh, T. & Akira, S. Autophagy in infection, inflammation and immunity. Nat Rev Immunol 13, 722–737 (2013).

113. G., B. B., et al. Inflammasome Components Coordinate Autophagy and Pyroptosis as Macrophage Responses to Infection. mBio 4, e00620–12 (2013).

114. Levine, B., Mizushima, N. & Virgin, H. W. Autophagy in immunity and inflammation. Nature 469, 323–335 (2011).

115. Siqueira, M. da S., Ribeiro, R. de M. & Travassos, L. H. Autophagy and Its Interaction With Intracellular Bacterial Pathogens. Front Immunol 9, 935 (2018).

116. Gordon, S. & Martinez, F. O. Alternative Activation of Macrophages: Mechanism and Functions. Immunity 32, 593–604 (2010).

117. Orecchioni, M., Ghosheh, Y., Pramod, A. B. & Ley, K. Macrophage Polarization: Different Gene Signatures in M1(LPS+) vs. Classically and M2(LPS–) vs. Alternatively Activated Macrophages. Front Immunol 10, (2019).

118. Viola, A., Munari, F., Sánchez-Rodríguez, R., Scolaro, T. & Castegna, A. The Metabolic Signature of Macrophage Responses. Front Immunol 10, (2019).

119. Torretta, S. et al. D-mannose suppresses macrophage IL-1β production. Nat Commun 11, 6343 (2020).

120. LeBleu, V. S. et al. PGC-1α mediates mitochondrial biogenesis and oxidative phosphorylation in cancer cells to promote metastasis. Nat Cell Biol 16, 992–1003 (2014).

121. Hong, E.-H. et al. Intratumoral injection of attenuated Salmonella vaccine can induce tumor microenvironmental shift from immune suppressive to immunogenic. Vaccine 31, 1377– 1384 (2013).

122. Duong, M. T.-Q., Qin, Y., You, S.-H. & Min, J.-J. Bacteria-cancer interactions: bacteria-based cancer therapy. Exp Mol Med 51, 1–15 (2019).

123. Sieow, B. F.-L., Wun, K. S., Yong, W. P., Hwang, I. Y. & Chang, M. W. Tweak to Treat: Reprograming Bacteria for Cancer Treatment. Trends Cancer 7, 447–464 (2021).

124. Pockley, A. G., Foulds, G. A., Oughton, J. A., Kerkvliet, N. I. & Multhoff, G. Immune cell phenotyping using flow cytometry. Curr Protoc Toxicol 66, 18 (2015).

125. Sun, T. & Altenbuchner, J. Characterization of a mannose utilization system in bacillus subtilis. J Bacteriol (2010) doi:10.1128/JB.01673-09.

126. Ryu, Y. H. et al. Differential immunostimulatory effects of Gram-positive bacteria due to their lipoteichoic acids. Int Immunopharm 9, 127–133 (2009).

127. Moreira, L. O. et al. The TLR2-MyD88-NOD2-RIPK2 signalling axis regulates a balanced pro-inflammatory and IL-10-mediated anti-inflammatory cytokine response to Gram-positive cell walls. Cell Microbiol 10, 2067–2077 (2008).

128. Holden, J. A. et al. Porphyromonas gingivalis lipopolysaccharide weakly activates M1 and M2 polarized mouse macrophages but induces inflammatory cytokines. Infect Immun 82, 4190–4203 (2014).

129. M., J. J., et al. Mannose receptor (CD206) activation in tumor-associated macrophages enhances adaptive and innate antitumor immune responses. Sci Transl Med 12, eaax6337 (2020).

130. García-González, G. et al. Triggering of protease-activated receptors (PARs) induces alternative M2 macrophage polarization with impaired plasticity. Mol Immunol 114, 278– 288 (2019).

131. Leber, N. et al. α-Mannosyl-Functionalized Cationic Nanohydrogel Particles for Targeted Gene Knockdown in Immunosuppressive Macrophages. Macromol Biosci 19, e1900162 (2019).

132. Stahl, P. D. & Ezekowitz, R. A. B. The mannose receptor is a pattern recognition receptor involved in host defense. Curr Opin Immunol 10, 50–55 (1998).

133. Murtaugh, M. P. & Foss, D. L. Inflammatory cytokines and antigen presenting cell activation. Vet Immunol Immunopathol 87, 109–121 (2002).

134. Zhang, X. & Mosser, D. M. Macrophage activation by endogenous danger signals. Journal of Pathology Preprint at 10.1002/path.2284 (2008).

135. Wang, L., Zhang, S., Wu, H., Rong, X. & Guo, J. M2b macrophage polarization and its roles in diseases. J Leukoc Biol 106, 345–358 (2019).

136. Graves, D. T. The potential role of chemokines and inflammatory cytokines in periodontal disease progression. Clinl Infect Dis 28, 482–490 (1999).

137. Domingues, C., AB da Cruz e Silva, O. & Henriques, A. Impact of cytokines and chemokines on Alzheimer’s disease neuropathological hallmarks. Curr Alzheimer Res 14, 870–882 (2017).

138. Chatzigeorgiou, A. et al. The pattern of inflammatory/anti-inflammatory cytokines and chemokines in type 1 diabetic patients over time. Ann Med 42, 426–438 (2010).

139. Tanaka, T. et al. Chemokines in tumor progression and metastasis. Cancer Sci 96, 317– 322 (2005).

140. Ben-Baruch, A. Host microenvironment in breast cancer development: inflammatory cells, cytokines and chemokines in breast cancer progression: reciprocal tumor– microenvironment interactions. Breast Can Res 5, 1–6 (2002).

141. Raman, D., Baugher, P. J., Thu, Y. M. & Richmond, A. Role of chemokines in tumor growth. Cancer Lett 256, 137–165 (2007).

142. Ramesh, G., MacLean, A. G. & Philipp, M. T. Cytokines and chemokines at the crossroads of neuroinflammation, neurodegeneration, and neuropathic pain. Mediators Inflamm 2013, 480739 (2013).

143. Benoit, M., Desnues, B. & Mege, J.-L. Macrophage Polarization in Bacterial Infections. J Immunol 181, 3733 LP – 3739 (2008).

144. Smith, T. D., Tse, M. J., Read, E. L. & Liu, W. F. Regulation of macrophage polarization and plasticity by complex activation signals. Integr Biol (Camb) 8, 946–955 (2016).

145. Rőszer, T. Understanding the Mysterious M2 Macrophage through Activation Markers and Effector Mechanisms. Mediators Inflamm 2015, 816460 (2015).

146. Martins, A., Han, J. & Kim, S. O. The multifaceted effects of granulocyte colony-stimulating factor in immunomodulation and potential roles in intestinal immune homeostasis. IUBMB Life 62, 611–617 (2010).

147. Becker, C. et al. Regulation of IL-12 p40 Promoter Activity in Primary Human Monocytes: Roles of NF-κB, CCAAT/Enhancer-Binding Protein β, and PU.1 and Identification of a Novel Repressor Element (GA-12) That Responds to IL-4 and Prostaglandin E2; J Immunol 167, 2608 LP – 2618 (2001).

148. Laha, D., Grant, R., Mishra, P. & Nilubol, N. The role of tumor necrosis factor in manipulating the immunological response of tumor microenvironment. Front Immunol 12, 656908 (2021).

149. Kenny, P. A., Lee, G. Y. & Bissell, M. J. Targeting the tumor microenvironment. Front Biosci 12, 3468 (2007).

150. Rosenzweig, J. M., Glenn, J. D., Calabresi, P. A. & Whartenby, K. A. KLF4 modulates expression of IL-6 in dendritic cells via both promoter activation and epigenetic modification. J Biol Chemy 288, 23868–23874 (2013).

151. Deshmane, S. L., Kremlev, S., Amini, S. & Sawaya, B. E. Monocyte chemoattractant protein-1 (MCP-1): an overview. J Interferon Cytokine Res 29, 313–326 (2009).

152. Sutcliffe, A. M. et al. Transcriptional regulation of monocyte chemotactic protein-1 release by endothelin-1 in human airway smooth muscle cells involves NF-kappaB and AP-1. Br J Pharmacol 157, 436–450 (2009).

153. Kanemaru, H., et al. Antitumor effect of Batf2 through IL-12 p40 up-regulation in tumor-associated macrophages. Proc Nat Acad Sci, USA 114, E7331 LP- E7340 (2017).

154. Chonov, D. C., Ignatova, M. M. K., Ananiev, J. R. & Gulubova, M. V. IL-6 Activities in the Tumour Microenvironment. Part 1. Open Access Maced J Med Sci 7, 2391–2398 (2019).

155. Hu, J. et al. Regulation of tumor immune suppression and cancer cell survival by CXCL1/2 elevation in glioblastoma multiforme. Sci Adv 7, eabc2511 (2021).

156. Burke, S. J. et al. NF-κB and STAT1 control CXCL1 and CXCL2 gene transcription. Am J Physiol Endocrinol Metab 306, E131–49 (2014).

157. Laoui, D. et al. Tumor-associated macrophages in breast cancer: distinct subsets, distinct functions. Int J Dev Biol 55, 861–867 (2011).

158. Bhavsar, I., Miller, C. S. & Al-Sabbagh, M. Macrophage Inflammatory Protein-1 Alpha (MIP-1 alpha)/CCL3: As a Biomarker. General Methods in Biomarker Research and their Applications 223–249 Preprint at 10.1007/978-94-007-7696-8_27 (2015).

159. Liu, D. et al. Comprehensive proteomics analysis reveals metabolic reprogramming of tumor-associated macrophages stimulated by the tumor microenvironment. J Proteome Res 16, 288–297 (2017).

160. Maduka, C. V et al. Glycolytic reprogramming underlies immune cell activation by polyethylene wear particles. Biomaterials Adv 152, 213495 (2023).

161. Maduka, C. V et al. Elevated oxidative phosphorylation is critical for immune cell activation by polyethylene wear particles. J Immunol Regen Med 19, 100069 (2023).

162. van Raam B. J., Sluiter, W., de Wit, E., Roos, D., Verhoeven, A. J., Kuijpers, T. W. Mitochondrial Membrane Potential in Human Neutrophils Is Maintained by Complex III Activity in the Absence of Supercomplex Organisation. PLoS One 3(4), e2013 (2008).

163. Mookerjee, S. A., Gerencser, A. A., Nicholls, D. G. & Brand, M. D. Quantifying intracellular rates of glycolytic and oxidative ATP production and consumption using extracellular flux measurements. J Biol Chem 292, 7189–7207 (2017).

164. Diskin, C. & Pålsson-McDermott, E. M. Metabolic Modulation in Macrophage Effector Function. Front Immunol 9, 270 (2018).

165. Rabold, K., Netea, M. G., Adema, G. J. & Netea-Maier, R. T. Cellular metabolism of tumor-associated macrophages–functional impact and consequences. FEBS Lett 591, 3022– 3041 (2017).

166. Wculek, S. K., Dunphy, G., Heras-Murillo, I., Mastrangelo, A. & Sancho, D. Metabolism of tissue macrophages in homeostasis and pathology. Cell Mol Immunol 19, 384–408 (2022).

167. Jawalagatti, V., Kirthika, P. & Lee, J. H. Targeting primary and metastatic tumor growth in an aggressive breast cancer by engineered tryptophan auxotrophic Salmonella Typhimurium. Mol Ther Oncolytics 25, 350 (2022).

168. Geng, Z. et al. Aptamer-assisted tumor localization of bacteria for enhanced biotherapy. Nat Commun 12, 6584 (2021).

169. Pan, H. et al. Engineered NIR light-responsive bacteria as anti-tumor agent for targeted and precise cancer therapy. Chem Eng J 426, 130842 (2021).

170. Zheng, D.-W. et al. Optically-controlled bacterial metabolite for cancer therapy. Nat Commun 9, 1680 (2018).

171. Daud, A. I. et al. Tumor immune profiling predicts response to anti–PD-1 therapy in human melanoma. J Clin Invest 126, 3447–3452 (2016).

172. Khalsa, J. K. et al. Immune phenotyping of diverse syngeneic murine brain tumors identifies immunologically distinct types. Nat Commun 11, 1–14 (2020).

173. DuPré, S. A., Redelman, D. & Hunter, K. W. J. The mouse mammary carcinoma 4T1: characterization of the cellular landscape of primary tumours and metastatic tumour foci. Int J Exp Pathol 88, 351–360 (2007).

174. Bronte, V. et al. Recommendations for myeloid-derived suppressor cell nomenclature and characterization standards. Nat Commun 7, 12150 (2016).

175. Gonzalez, P. S. et al. Mannose impairs tumour growth and enhances chemotherapy. Nature 563, 719–723 (2018).

176. Liu, Q., Li, X., Zhang, H. & Li, H. Mannose Attenuates Colitis-Associated Colorectal Tumorigenesis by Targeting Tumor-Associated Macrophages. J Cancer Prev 27, 31 (2022).

177. Whiteside, T. L. What are regulatory T cells (Treg) regulating in cancer and why? in Semin Cancer Biol vol. 22 327–334 (Elsevier, 2012).

178. Rech, A. J. et al. CD25 blockade depletes and selectively reprograms regulatory T cells in concert with immunotherapy in cancer patients. Sci Transl Med 4, 134ra62–134ra62 (2012).

179. Sharma, S. et al. Tumor cyclooxygenase-2/prostaglandin E2–dependent promotion of FOXP3 expression and CD4+ CD25+ T regulatory cell activities in lung cancer. Cancer Res 65, 5211–5220 (2005).

180. Lopez-Cabrera, M. et al. Molecular cloning, expression, and chromosomal localization of the human earliest lymphocyte activation antigen AIM/CD69, a new member of the C-type animal lectin superfamily of signal-transmitting receptors. J Exp Med 178, 537–547 (1993).

181. Fontenot, J. D., Gavin, M. A. & Rudensky, A. Y. Foxp3 programs the development and function of CD4+ CD25+ regulatory T cells. Nat Immunol 4, 330–336 (2003).

182. Lio, C.-W. J. & Hsieh, C.-S. A two-step process for thymic regulatory T cell development. Immunity 28, 100–111 (2008).

183. Vang, K. B. et al. IL-2,-7, and-15, but not thymic stromal lymphopoeitin, redundantly govern CD4+ Foxp3+ regulatory T cell development. J Immunol 181, 3285–3290 (2008).

184. Klein Geltink, R. I., Kyle, R. L. & Pearce, E. L. Unraveling the complex interplay between T cell metabolism and function. Annu Rev Immunol 36, 461–488 (2018).

185. Rokop, M. E., Auchtung, J. M. & Grossman, A. D. Control of DNA replication initiation by recruitment of an essential initiation protein to the membrane of Bacillus subtilis. Mol Microbiol 52, 1757–67 (2004).

186. Harwood, C. R. & Cutting, S. M. Molecular biological methods for Bacillus. Preprint at (1990).

187. Francis, K. P. et al. Visualizing pneumococcal infections in the lungs of live mice using bioluminescent Streptococcus pneumoniae transformed with a novel gram-positive lux transposon. Infect Immun (2001) doi:10.1128/IAI.69.5.3350-3358.2001.

188. Gonçalves, R. & Mosser, D. M. The Isolation and Characterization of Murine Macrophages. Curr Protoc Immunol 111, 14.1.1–14.1.16 (2015).

189. Mills, E. L. et al. Succinate Dehydrogenase Supports Metabolic Repurposing of Mitochondria to Drive Inflammatory Macrophages. Cell 167, 457–470.e13 (2016).

190. Zhang, G.-L., Zhang, Y., Cao, K.-X. & Wang, X.-M. Orthotopic Injection of Breast Cancer Cells into the Mice Mammary Fat Pad. JoVE e58604 (2019) doi:doi:10.3791/58604.

191. Liu, X. et al. CD47 blockade triggers T cell–mediated destruction of immunogenic tumors. Nat Med 21, 1209–1215 (2015).

