## Extended data figures for "Engineered endosymbionts that modulate primary macrophage function and attenuate tumor growth by shifting the tumor microenvironment"

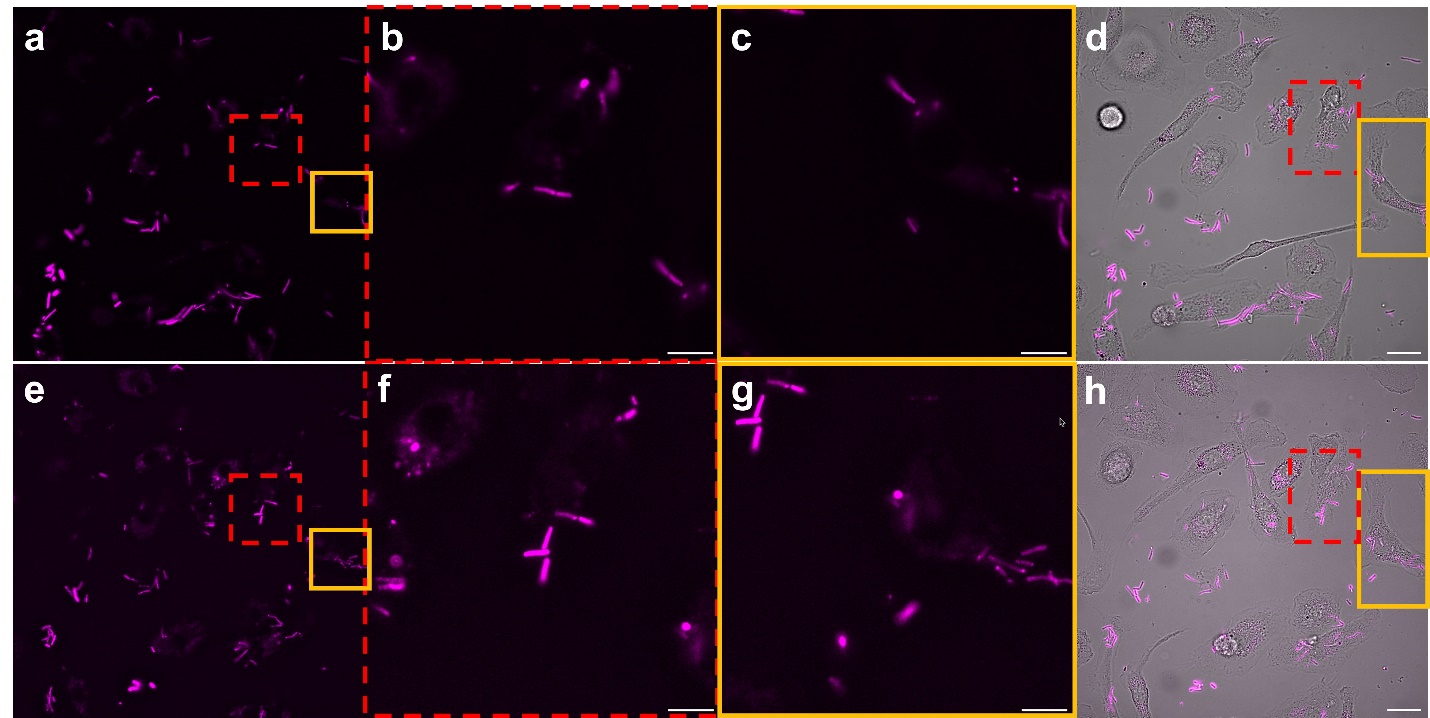

Extended Fig. 1. Live cell imaging of replicating *B. subtilis* LLO inside live BMDMs

Live cell microscopy revealed the LLO strain replicating in multiple host cells by comparing images at 3 h (top images, a-d) and 4.5 h (bottom images, e-f) post-bacterial addition. BMDMs were visualized in brightfield (d, h), and the LLO strain using fluorescence (magenta); zoomed images (b-d and f-g) reveal bacteria (magenta LLO-expressing strain of *B. subtilis*) replicating in the cytoplasm and the regions at higher resolution are marked on low resolution images with red dashed box or a yellow solid box. Scale bars = 3 µm for zoomed images (b-c and f-g) and scale bars = 20 µm for not zoomed images are in overlay images (d, h).

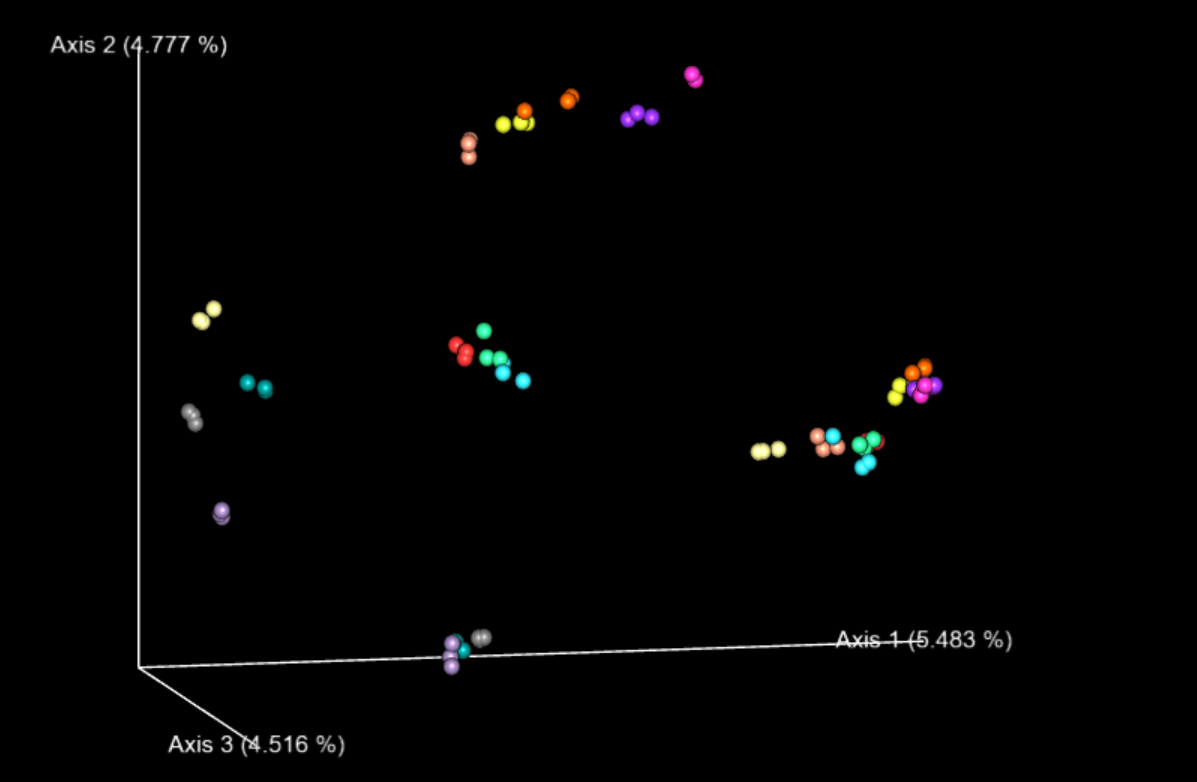

**Extended Fig. 2. Genome-wide gene expression shifts in BMDMs after response to engineered *B. subtilis* LLO strains and controls**

QIIME 2 Emperor Plot visualizes the shifts in genome-wide gene expression after BMDMs were untreated (gray), treated with LPS (salmon), mannose (lavender), LPS and IFN-γ (pink), IL-4 and IL-13 (teal), LLO strain without IPTG (no IPTG, sand), LLO strain with and without mannose (LLO -mannose, orange; LLO +mannose, neon green), LLO-*SK* with and without mannose (LLO-*SK* -mannose, purple; LLO-*SK­* +mannose, light blue) and LLO-*KG* with and without mannose (LLO-*KG* -mannose, yellow; LLO-*KG* +mannose, red) at 12 and 24 h post-initial treatment. The 24 h treatments are positioned in the middle, left and upper side of the plot while the 12 h are on the bottom and right side of the plot in this 2D view. The 3D visualization can be accessed through QIIME 2 view by the process explained in the GitHub.

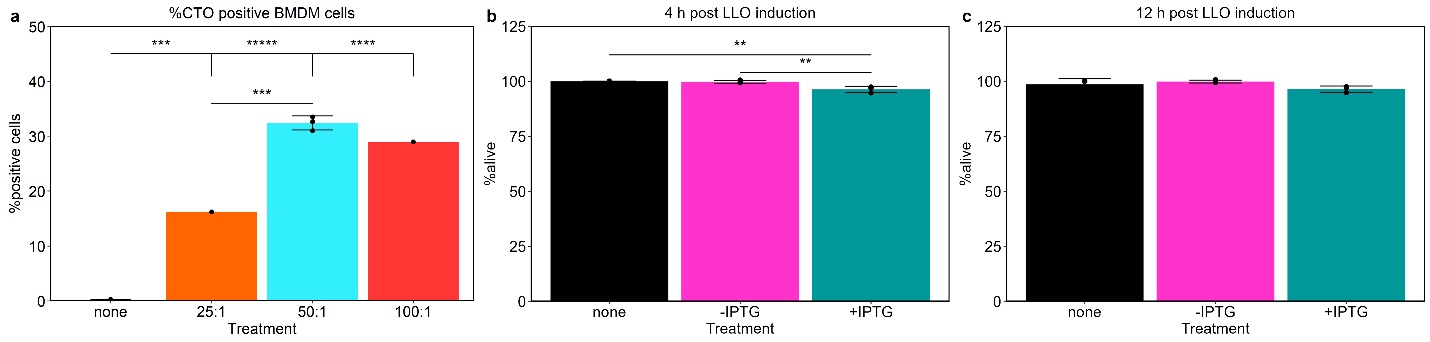

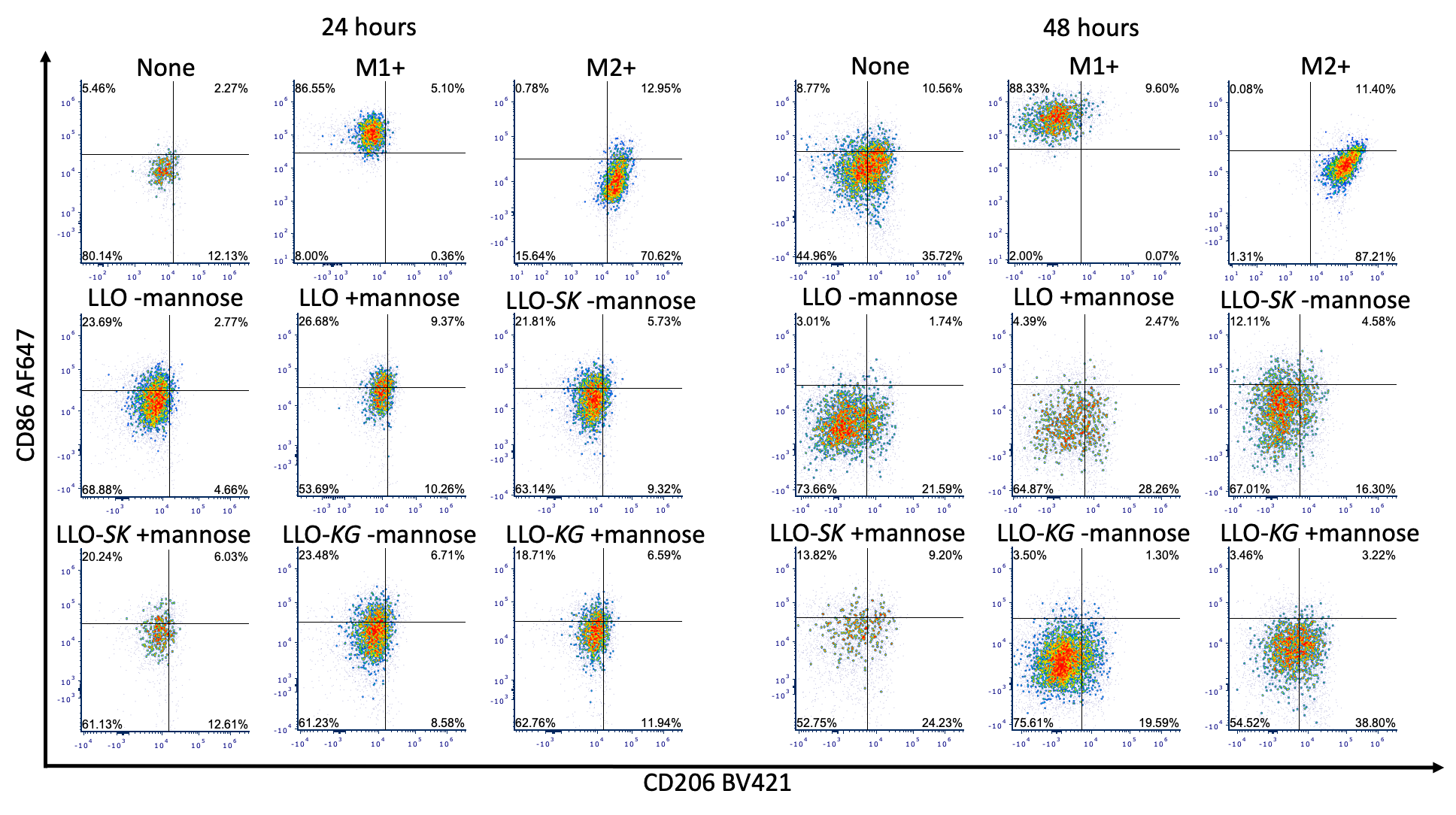

Extended Fig. 3. Uptake of *B. subtilis* LLO by BMDM and assessment of BMDM viability after uptake

BMDMs were co-incubated with *B. subtilis* LLO strain (+IPTG) which were stained with CellTracker Orange (CTO) CMRA Dye. CTO-stained LLO were added at different MOIs and incubated for 4 h to determine number of cells with bacteria (a). BMDMs were analyzed for viability using flow cytometry at 4 and 12 h at a 50:1 MOI (b-c). Experiments were performed with three biological replicates (n=3) except for 25:1 and 100:1 MOI in uptake experiment (a) which were performed with one biological replicate (n=1) to correlate with live cell data trends. Data is mean ± SD; **p<0.01, ***p<0.001, ****p<0.0001, *****p<0.00001.

**Extended Fig. 4. Changes in the levels of surface markers on BMDMs, by flow** **cytometry, after various treatments with *B. subtilis***

Flow cytometry dot plots of CD86 AF647 and CD206 BV421 surface markers after BMDMs were untreated (none), treated with LPS and IFN-γ (M1+), IL-4 and IL-13 (M2+), LLO strain with and without mannose (LLO -mannose, LLO +mannose), LLO-*SK* with and without mannose (LLO-*SK* -mannose, LLO-*SK­* +mannose) and LLO-*KG* with and without mannose (LLO-*KG* -mannose, LLO-*KG* +mannose) at 24 and 48 h post-initial treatment. IPTG was added to all bacterial treatments. Percentages expressed in quadrants are from live, CD11b+/F4/80+ population.

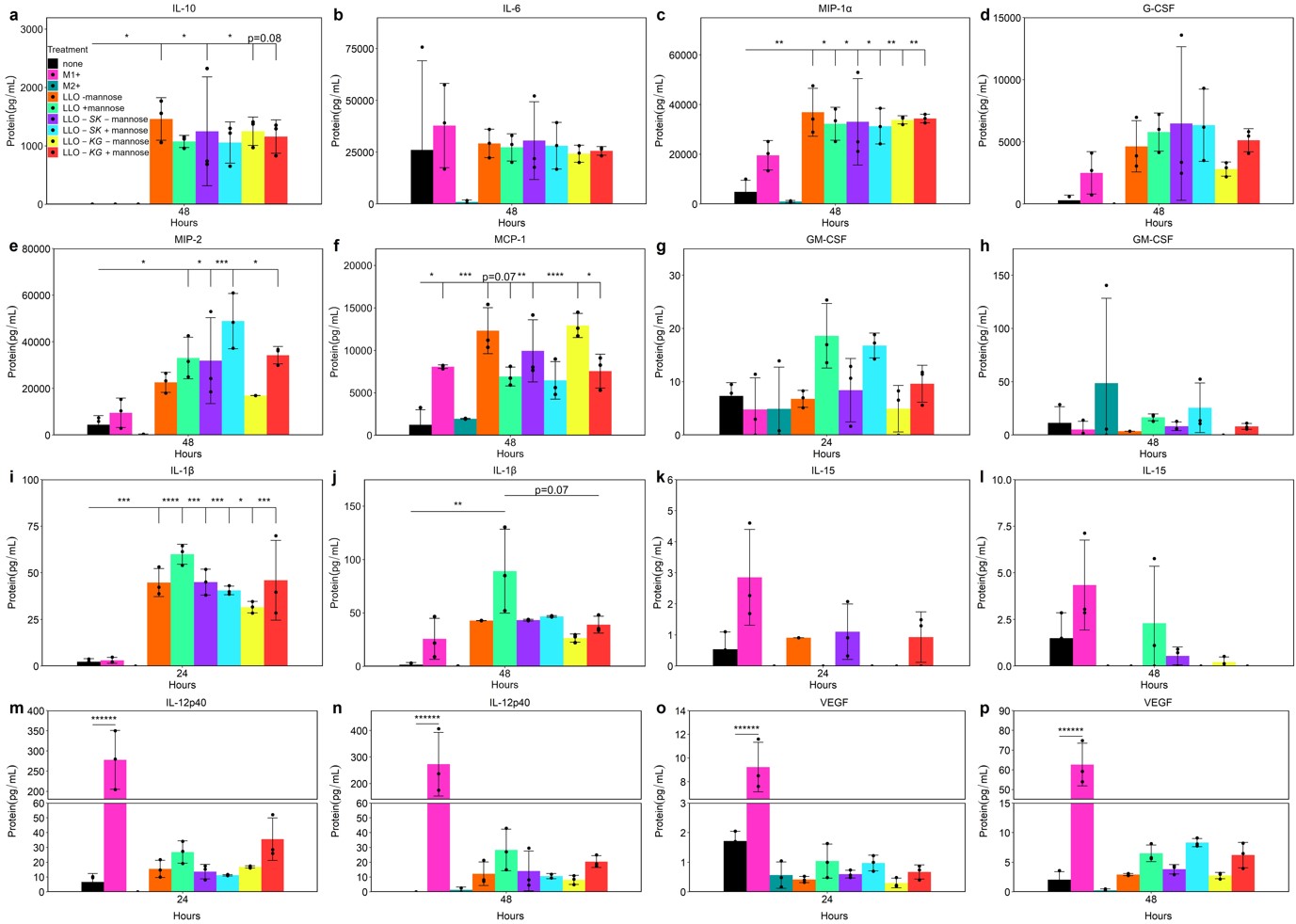

**Extended Fig. 5. Additional profiling of BMDM cytokine and chemokine production after exposure to engineered *B. subtilis* LLO strains**

Cytokine and chemokine protein concentrations were quantified after BMDMs cells were untreated (none), treated with LPS and IFN-γ (M1+), IL-4 and IL-13 (M2+), LLO strain with and without mannose (LLO -mannose, LLO +mannose), LLO-*SK* with and without mannose (LLO-*SK* -mannose, LLO-*SK­* +mannose) and LLO-*KG* with and without mannose (LLO-*KG* -mannose, LLO-*KG* +mannose) at 24 h and 48 h post-initial treatment. IPTG was added to all bacterial treatments. Data is mean ± SD from n=3 biological replicates; *p<0.05, **p<0.01, ***p<0.001, ****p<0.0001, *****p<0.00001, ******p<0.000001.

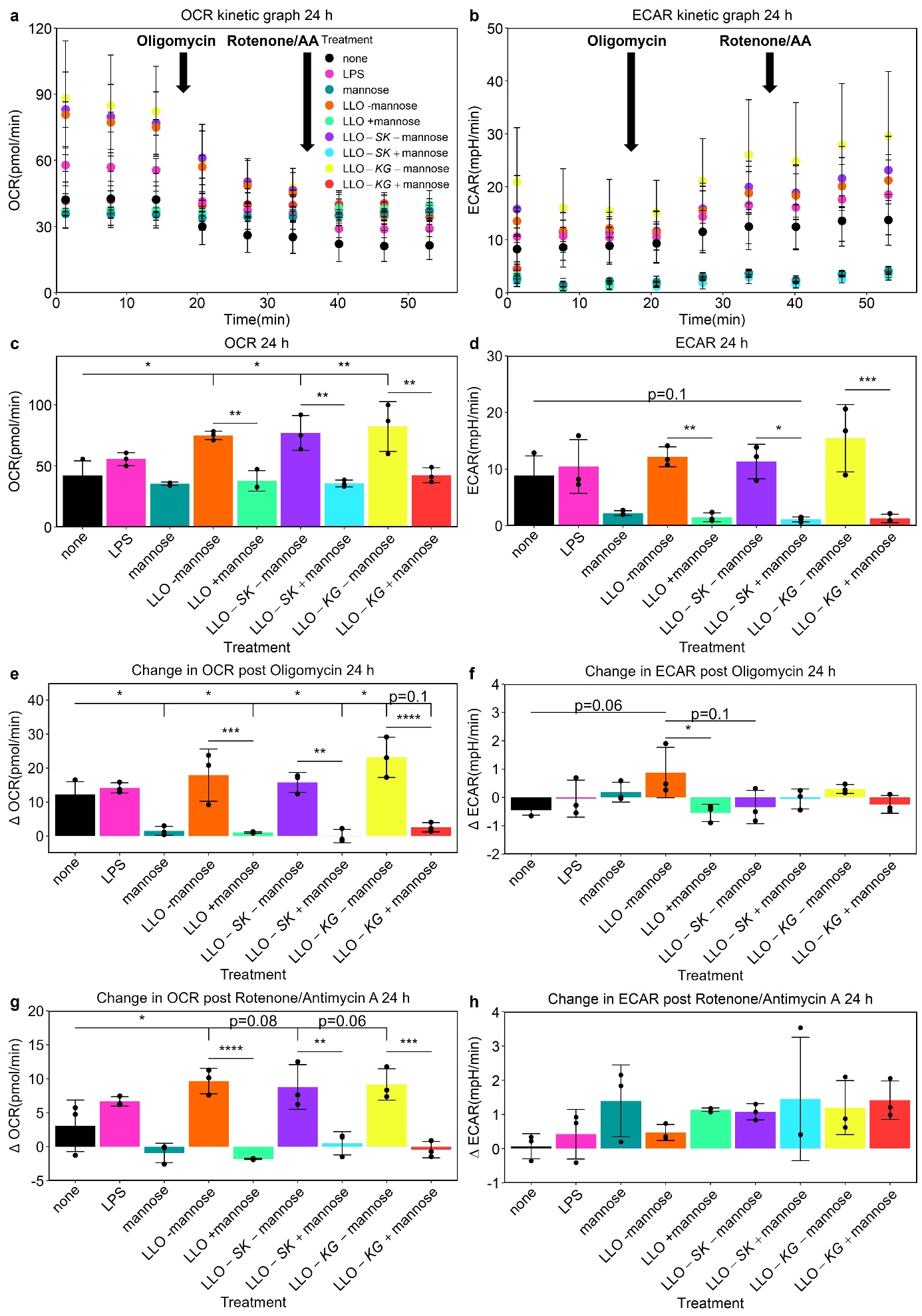

Extended Fig. 6. Patterns of functional metabolism for BMDMs 24 h after exposure to engineered *B. subtilis* LLO variants

OCR and ECAR were measured before and after electron transport chain inhibitors, Oligomycin and Rotenone/antimycin A (AA), were added at points indicated on the kinetic plots (a, b). OCR and ECAR quantification at the third measurement before addition of inhibitors were plotted (c, d). Further analysis was performed to quantify changes in OCR (ΔOCR) and ECAR (ΔECAR) after inhibitors were added (e-h). BMDMs were untreated (none), treated with LPS, mannose, LLO strain with and without mannose (LLO -mannose, LLO +mannose), LLO-*SK* with and without mannose (LLO-*SK* -mannose, LLO-*SK­* +mannose) and LLO-*KG* with and without mannose (LLO-*KG* -mannose, LLO-*KG* +mannose). IPTG was added to all bacterial treatments. Data is mean ± SD from n=3 biological replicates; *p<0.05, **p<0.01, ***p<0.001, ****p<0.0001.

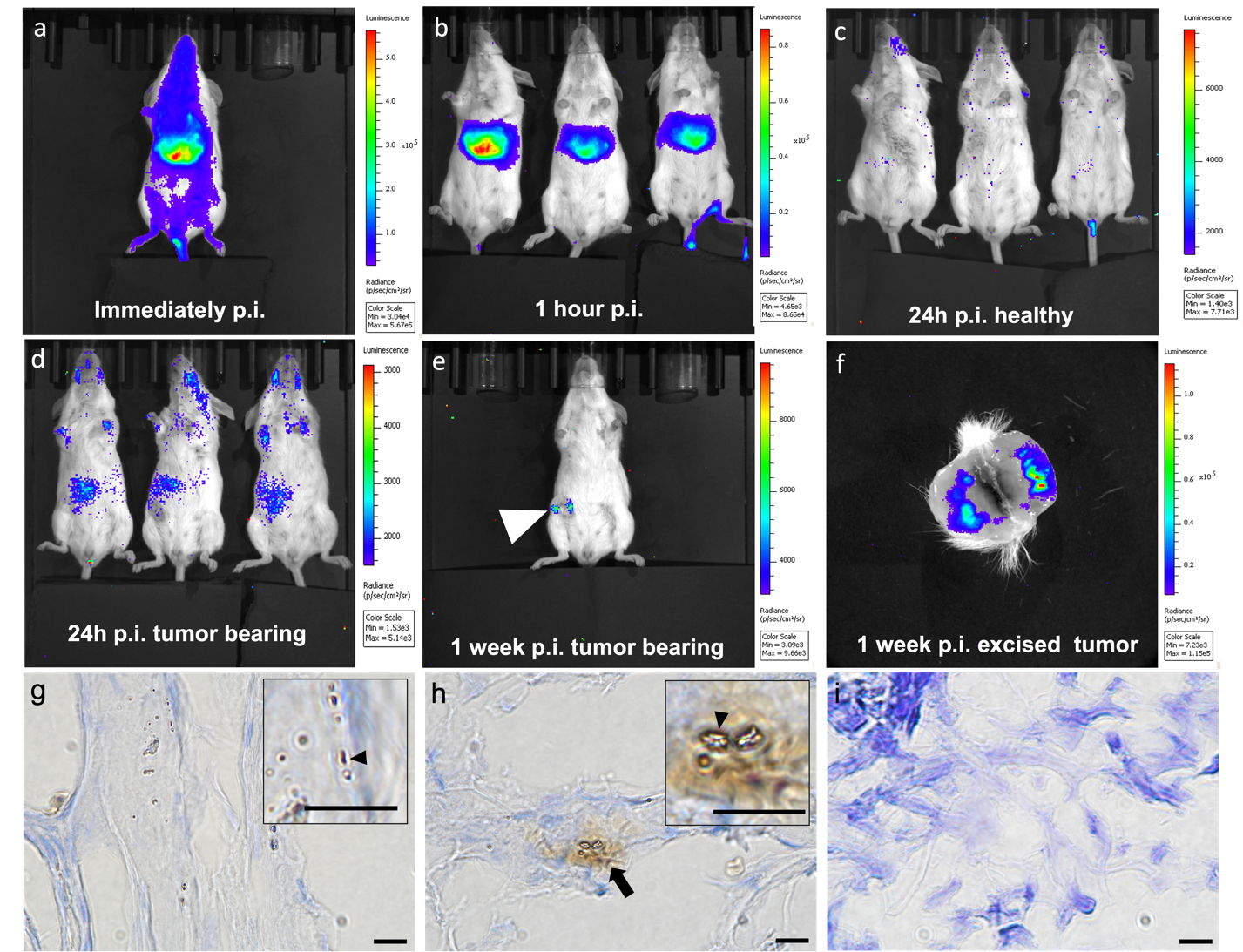

**Extended Fig. 7*. In vivo* bioluminescence imaging of the LLO-*lux* strain in mice after intravenous injection into representative healthy and tumor bearing mice and end point tumor histology.**

Representative healthy and 4T1 orthotopic tumor bearing mice were injected with 10^8^ LLO-*lux* intravenously and followed using *in vivo* bioluminescence imaging (BLI) to monitor LLO-*lux* location and viability. Representative images are shown immediately after injection (a), then at 1 h post-injection (p.i.; B), 24 h p.i. (d) and 1-week p.i. (e, f). Healthy and tumor bearing mice showed the same trends immediately after injection and 1 h p.i. However, by 24 h, all bacterial signal is cleared from healthy mice (c). Bacteria were shown to persist throughout the tumor 1-week p.i. which was identified within the tumor in an intact animal (e) and in more detail, within an excised tumor which was cut in half (f). Immunohistochemistry identified rod-shaped bacteria (black arrowheads) in tumors which were injected intravenous (IV) with the bacteriotherapeutic. These were present both in CD11b- cells (g) and CD11b+ cells (h, brown). There were no bacteria observed in tumors of mice which did not receive bacteria injection (i). Scale bars = 20 µm.

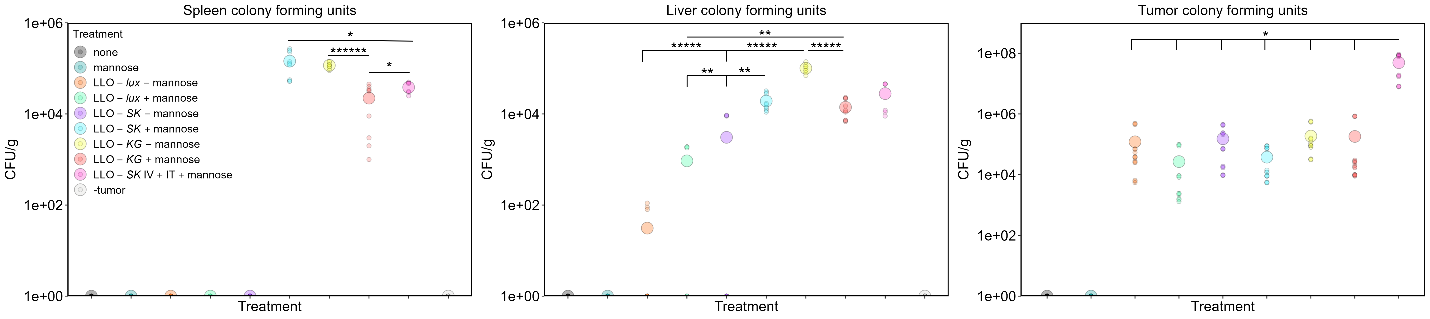

**Extended Fig. 8. Colony forming units from tumors and relevant organs**

Colony forming units (CFU) were calculated and normalized to mass (g) of spleens, livers and tumors. Groups were untreated (none), treated with mannose, LLO-*lux* strain injected IV with and without mannose (LLO-*lux* -mannose, LLO-*lux* +mannose), LLO-*SK* injected IV with and without mannose (LLO-*SK* -mannose, LLO-*SK­* +mannose), LLO-*KG* injected IV with and without mannose (LLO-*KG* -mannose, LLO-*KG* +mannose), LLO-*SK* injected IV and IT with mannose (LLO-*SK­* IV+IT +mannose) and no tumors (-tumor). IPTG was added to all bacterial treatments. Data is a scatter plot of all individual values with large point representing the mean from n=3 spleens/livers and n=5 tumors (3 halves and 2 whole) that were plated in triplicate from each treatment.

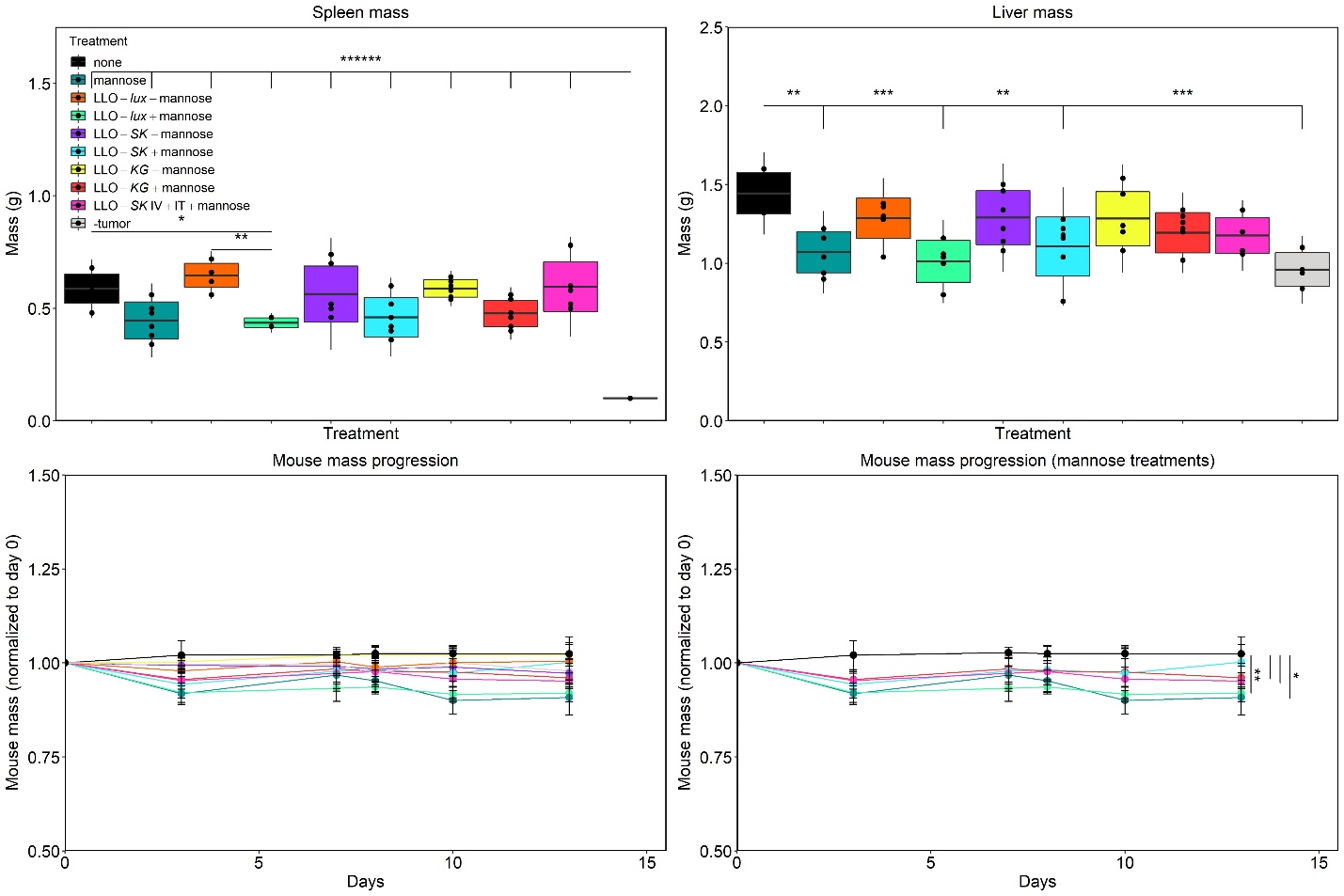

**Extended Fig. 9. Changes in body weight (mass progression) of mice and final mass of organs found to contain bacteria**

The mass of spleens and liver (top panels) and mice (lower panels) after the following treatments: untreated (none), treated with mannose, LLO-*lux* strain injected IV with and without mannose (LLO-*lux* -mannose, LLO-*lux* +mannose), LLO-*SK* injected IV with and without mannose (LLO-*SK* -mannose, LLO-*SK­* +mannose), LLO-*KG* injected IV with and without mannose (LLO-*KG* -mannose, LLO-*KG* +mannose), LLO-*SK* injected IV and IT with mannose (LLO-*SK­* IV+IT +mannose) and healthy mice injected with LLO-*lux* (-tumor). IPTG was added to all bacterial treatments. Data is mean ± SD for box and two times ± SD for whiskers from n=6 mice (tumor bearing) and n=4 mice (-tumor) for upper panels and mean ± SD from same mice for lower panels.

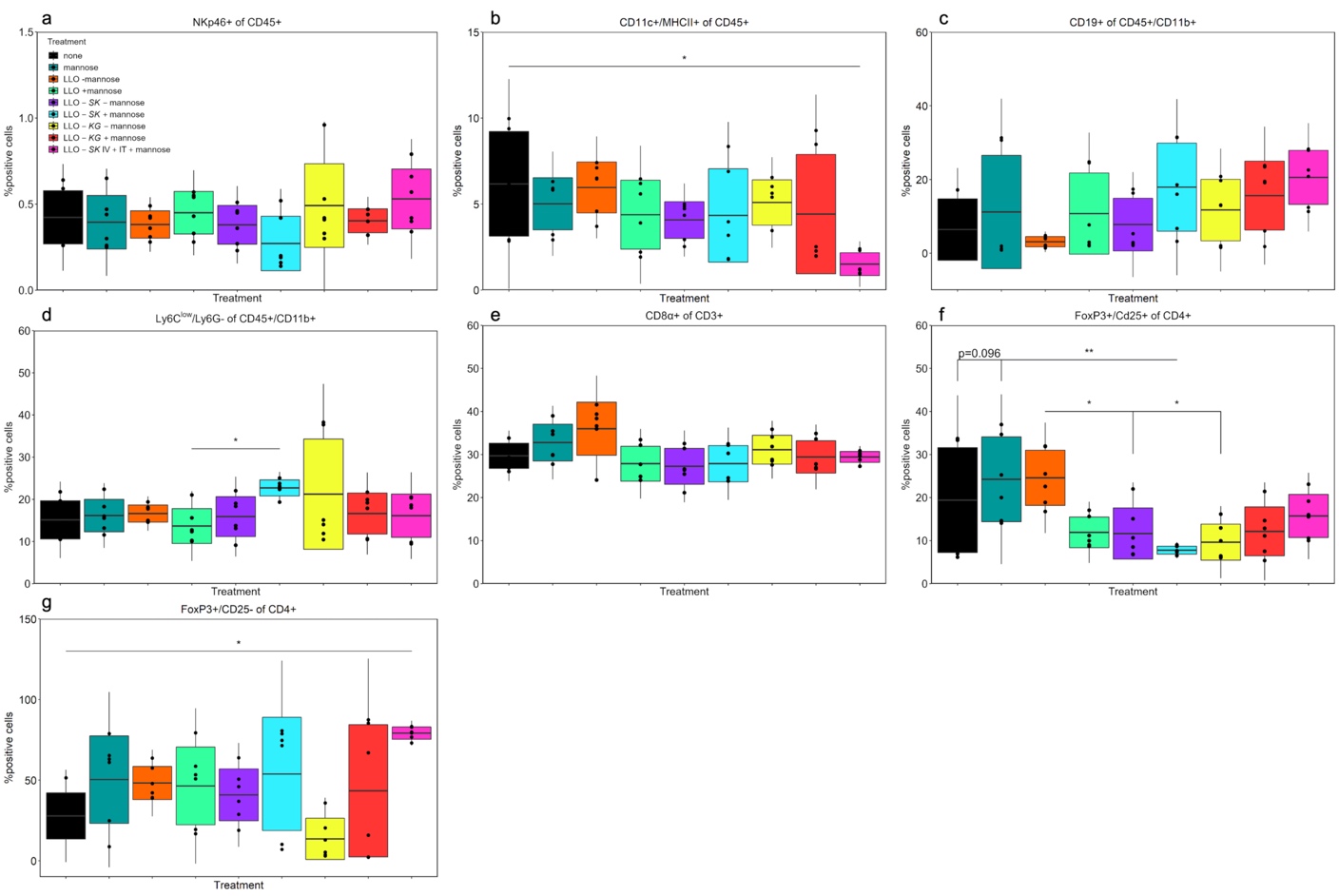

**Extended Fig. 10. Immunophenotyping of 4T1 tumors after treatment**

Immune cell populations were analyzed using flow cytometry, from digested tumors from all treatment groups: untreated (none), treated with mannose, LLO-*lux* strain injected IV with and without mannose (LLO-*lux* -mannose, LLO-*lux* +mannose), LLO-*SK* injected IV with and without mannose (LLO-*SK* -mannose, LLO-*SK­* +mannose), LLO-*KG* injected IV with and without mannose (LLO-*KG* -mannose, LLO-*KG* +mannose) and LLO-*SK* injected IV and IT with mannose (LLO-*SK­* IV+IT +mannose). IPTG was added to all bacterial treatments. Populations analyzed are NK cells (NKp46^+^ of CD45^+^), dendritic cells (CD11c^+^/MHCII^+^ of CD45^+^), B cells (CD19^+^ of CD45^+^/CD11b^+^), tumor associated macrophages (TAMs; Ly6C^low^/Ly6G^-^ of CD45^+^/CD11b^+^), cytotoxic T cells (CD8α^+^ of CD3^+^), T regulatory cells (Treg; FoxP3^+^/CD25^+^ of CD4^+^) and an alternative Treg subset (FoxP3^+^/CD25^-^ of CD4^+^). Data is mean ± SD for box and two times ± SD for whiskers from n=3 tumors in technical duplicates for each treatment; *p<0.05, **p<0.01.
