## Supplemental data for "Engineered endosymbionts that modulate primary macrophage function and attenuate tumor growth by shifting the tumor microenvironment"

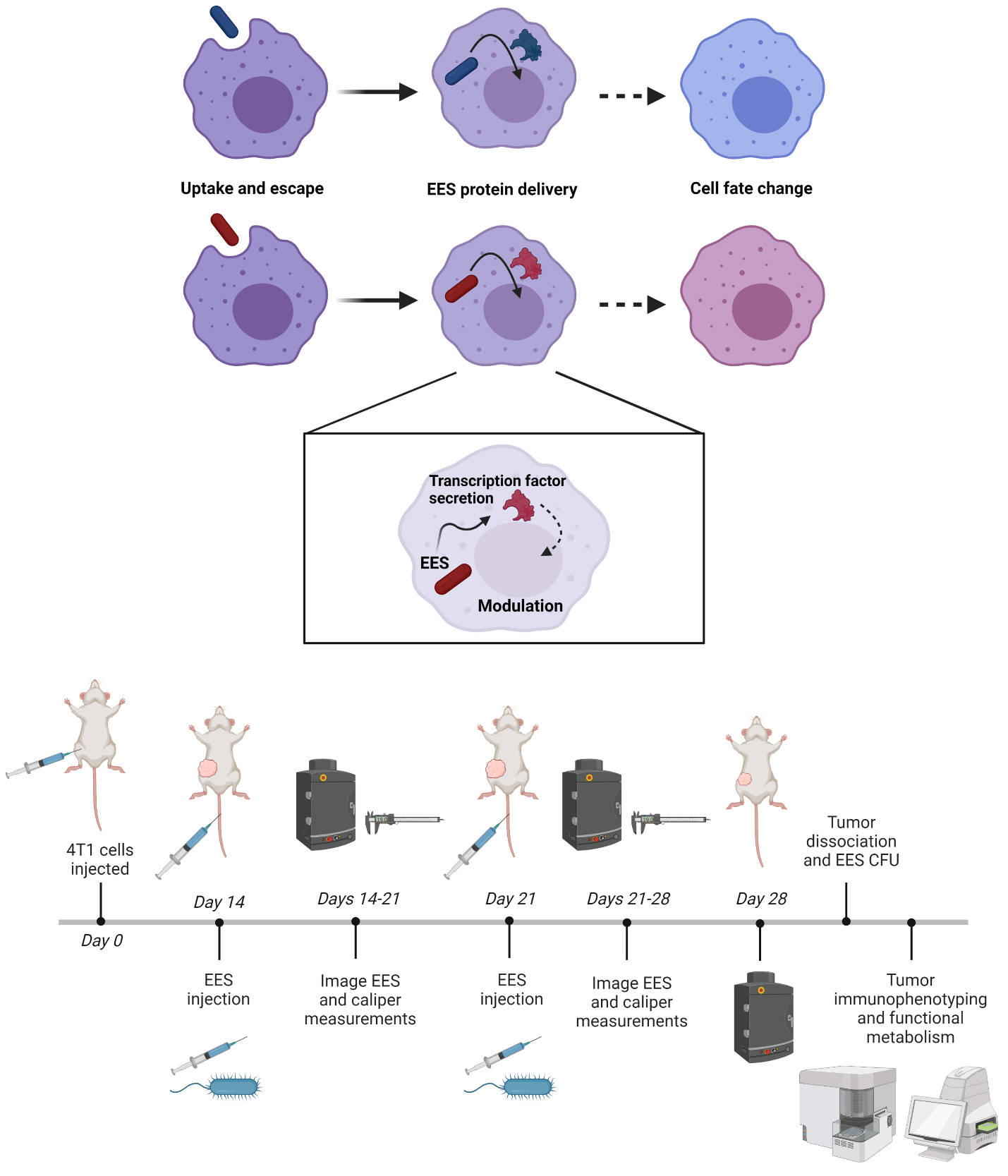

**Supplementary Fig. 1. The design of EES to modulate macrophage function and demonstration of EES-mediated modulation of the tumor microenvironment**

The EES are taken up by macrophages and escape into the cytoplasm to deliver TFs to modulate host cell function. To demonstrate utility redirecting the TME, we used an orthotopic 4T1 breast cancer model in mice and evaluated the localization of EES and effects on the TME. The EES localization and persistence were tracked using an In Vivo Imaging System (IVIS), and tumor growth was measured by calipers. Tumors were characterized by immunophenotyping (flow cytometry), functional metabolism (Seahorse real-time metabolic assays) and EES colony forming units (CFU).

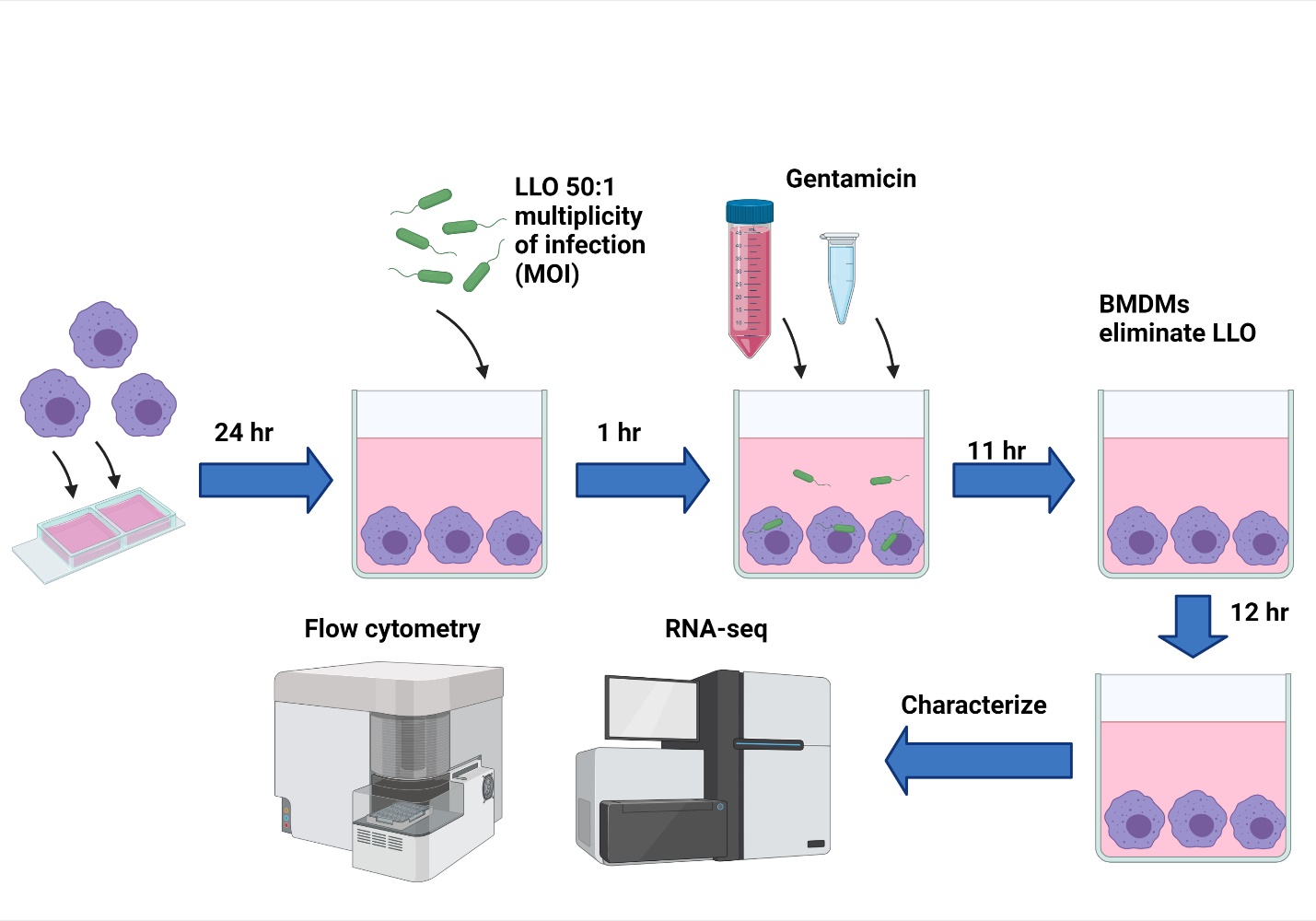

Supplementary Fig. 2. Diagram of method to deliver *B. subtilis* LLO strains to BMDMs and analyze interactions

*B. subtilis* LLO were co-incubated with host BMDMs according to the timeline shown and analyzed. Bacteria were incubated with BMDMs for 1 h then gentamicin was added to eliminate extracellular bacteria. After 11 additional hours of incubation, BMDMs were observed to eliminate all intracellular bacteria. Analysis was performed at multiple time intervals but in most cases, incubation was continued for an additional 12 h to determine impact on host cells by flow cytometry and RNA-sequencing.

Supplementary Table 1. Quantification of shifts in functional metabolism shifts between glycolysis and oxidative phosphorylation (as percent) and resulting differences in ATP production rates at 12 h

Quantification of shifts between glycolysis and oxidative phosphorylation (shown in percent) and resulting impact on ATP production (by two measures to create an index) when BMDMs were untreated (none), treated with LPS, mannose, LLO strain with and without mannose (LLO -mannose, LLO +mannose), LLO-*SK* with and without mannose (LLO-*SK* -mannose, LLO-*SK­* +mannose) and LLO-*KG* with and without mannose (LLO-*KG* -mannose, LLO-*KG* +mannose) at 12 h post-addition of treatments.

|  | **Basal Rates (Average) 12 h** | | | | | | | | | | | |
| --- | --- | --- | --- | --- | --- | --- | --- | --- | --- | --- | --- | --- |
|  | **glycoATP Production Rate(pmol/min)** | | **mitoATP Production Rate (pmol/min)** | | **Total ATP Production Rate(pmol/min)** | | **XF ATP Rate Index** | | **% Glycolysis** | | **% Oxidative Phosphorylation** | |
| **Groups** | **Average** | **StDev** | **Average** | **StDev** | **Average** | **StDev** | **Average** | **StDev** | **Average** | **StDev** | **Average** | **StDev** |
| Untreated  (none) | 7.4 | 3.2 | 28.3 | 11.2 | 35.8 | 13.7 | 4.1 | 1.6 | 20.9 | 6.5 | 79.1 | 6.5 |
| LPS 100ng/ mL | 38.7 | 7.7 | 67.3 | 18.0 | 106.0 | 25.7 | 1.7 | 0.1 | 36.8 | 1.5 | 63.2 | 1.5 |
| Mannose | 2.9 | 8.6 | 6.0 | 1.1 | 8.9 | 9.8 | -2.596 | 3.810 | -46.335 | 127.003 | 146.3 | 127.0 |
| LLO | 39.1 | 17.9 | 90.1 | 18.5 | 129.2 | 36.0 | 2.5 | 0.8 | 29.3 | 6.0 | 70.7 | 6.0 |
| LLO +mannose | 48.1 | 6.9 | 88.5 | 17.9 | 136.6 | 24.2 | 1.8 | 0.2 | 35.4 | 2.6 | 64.6 | 2.6 |
| LLO-SK | 31.9 | 15.5 | 87.2 | 27.2 | 119.1 | 42.2 | 2.9 | 0.5 | 26.1 | 3.7 | 73.9 | 3.7 |
| LLO-SK +mannose | 28.7 | 12.0 | 66.9 | 15.1 | 95.7 | 25.3 | 2.6 | 0.9 | 29.2 | 7.1 | 70.8 | 7.1 |
| LLO-KG | 60.3 | 17.1 | 125.3 | 34.0 | 185.6 | 51.0 | 2.1 | 0.1 | 32.5 | 1.0 | 67.5 | 1.0 |
| LLO-KG +mannose | 59.4 | 20.5 | 105.2 | 43.3 | 164.6 | 63.6 | 1.8 | 0.2 | 36.4 | 2.2 | 63.6 | 2.2 |

Supplementary Table 2. Quantification of shifts in functional metabolism between glycolysis and oxidative phosphorylation (as percent) and resulting differences in ATP production rates at 24 h

Quantification of shifts between glycolysis and oxidative phosphorylation (shown in percent) and resulting impact on ATP production (by two measures to create an index) when BMDMs were untreated (none), treated with LPS, mannose, LLO strain with and without mannose (LLO -mannose, LLO +mannose), LLO-*SK* with and without mannose (LLO-*SK* -mannose, LLO-*SK­* +mannose) and LLO-*KG* with and without mannose (LLO-*KG* -mannose, LLO-*KG* +mannose) at 24 h post-addition of treatments.

|  | **Basal Rates (Average) 24 h** | | | | | | | | | | | |
| --- | --- | --- | --- | --- | --- | --- | --- | --- | --- | --- | --- | --- |
|  | **glycoATP Production Rate(pmol/min)** | | **mitoATP Production Rate (pmol/min)** | | **Total ATP Production Rate(pmol/min)** | | **XF ATP Rate Index** | | **% Glycolysis** | | **% Oxidative Phosphorylation** | |
| **Groups** | **Average** | **StDev** | **Average** | **StDev** | **Average** | **StDev** | **Average** | **StDev** | **Average** | **StDev** | **Average** | **StDev** |
| Untreated | 68.0 | 27.3 | 93.2 | 31.3 | 161.1 | 56.7 | 1.4 | 0.3 | 42.0 | 5.7 | 58.0 | 5.7 |
| LPS 100ng/ mL | 79.1 | 42.2 | 109.2 | 11.7 | 188.3 | 53.3 | 1.6 | 0.6 | 40.2 | 9.7 | 59.8 | 9.7 |
| Mannose | 20.1 | 4.1 | 8.3 | 7.4 | 28.5 | 11.3 | 0.38 | 0.27 | 74.2 | 14.4 | 25.8 | 14.4 |
| LLO | 86.3 | 15.0 | 165.0 | 13.0 | 251.4 | 26.5 | 1.9 | 0.2 | 34.2 | 2.6 | 65.8 | 2.6 |
| LLO +mannose | 13.3 | 6.8 | 5.9 | 1.2 | 19.2 | 6.9 | 0.52 | 0.28 | 67.3 | 11.6 | 32.7 | 11.6 |
| LLO-SK | 78.9 | 22.2 | 167.0 | 35.2 | 246.0 | 55.3 | 2.2 | 0.4 | 31.8 | 3.6 | 68.2 | 3.6 |
| LLO-SK +mannose | 9.4 | 3.9 | 5.3 | 6.1 | 14.7 | 10.0 | 0.44 | 0.45 | 74.3 | 23.8 | 25.7 | 23.8 |
| LLO-KG | 113.0 | 45.1 | 190.5 | 64.2 | 303.5 | 109.3 | 1.7 | 0.2 | 36.8 | 2.0 | 63.2 | 2.0 |
| LLO-KG +mannose | 10.5 | 6.0 | 14.6 | 8.9 | 25.1 | 14.8 | 1.4 | 0.2 | 42.0 | 4.3 | 58.0 | 4.3 |

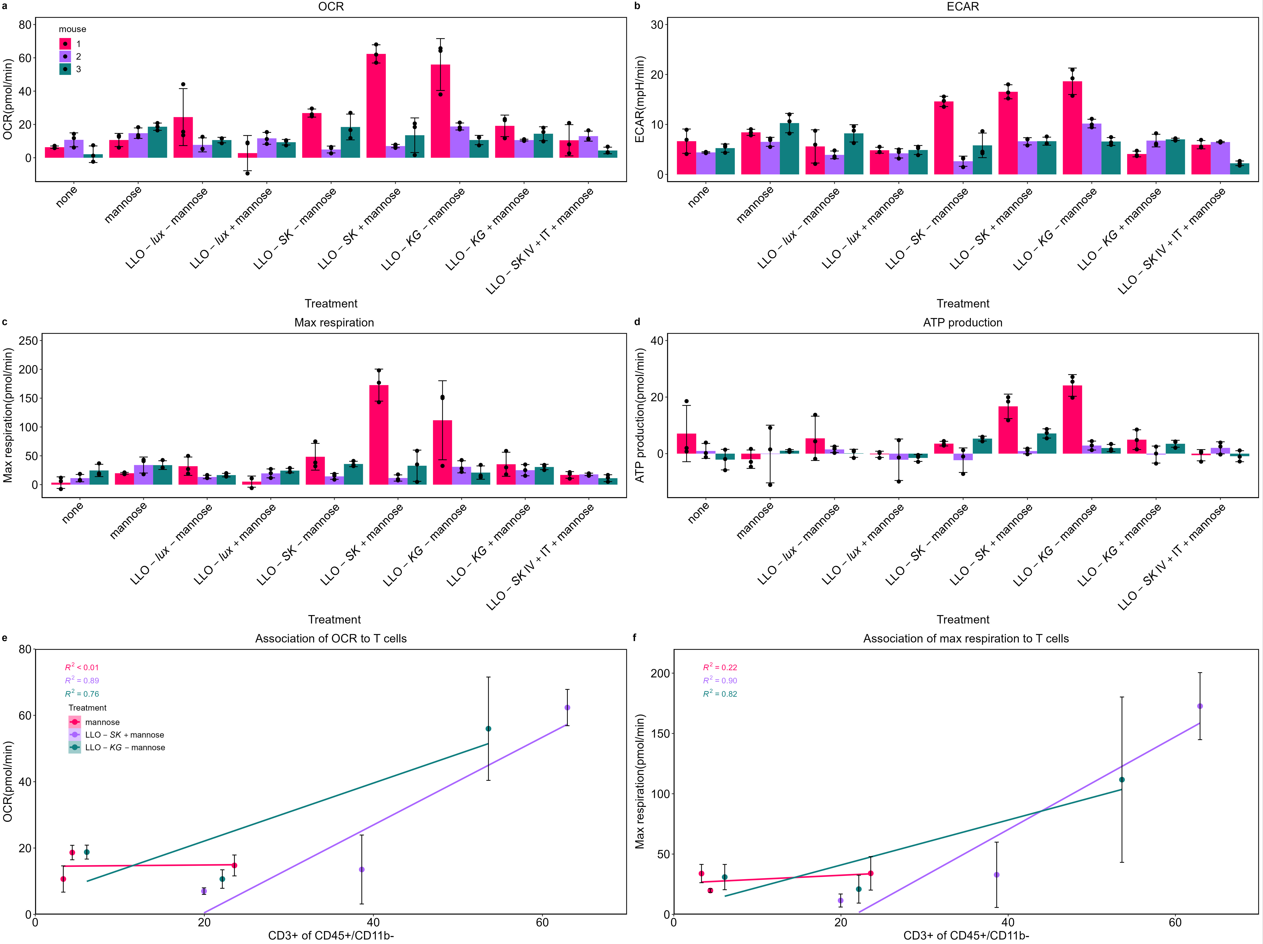

Supplementary Fig. 3. Functional metabolism of cells in the tumor microenvironment after treatment

OCR, ECAR, max respiration and ATP production were measured from cells released from resected tumors from treated mice (n=3 mice, n=3 technical replicates). Individual measurements from the tumors from each mouse were plotted and grouped into treatment schemes (a-d). Example associations between OCR or Max respiration measurements and CD3+ T cells (taken from the same tumors, split in half for each analysis) are plotted with the two treatment groups, with the highest association, and one with a lower association shown. R^2^ values are noted for each treatment group. Data is mean ± SD.

Supplementary Table 3. Summary of liver histopathology

The livers of tumor-bearing mice in five treatment groups [1 (untreated), 2 (mannose-treated), 4 (LLO +mannose), 6 (LLO-*SK* +mannose) and 8 (LLO-*KG* +mannose)] were excised, fixed, sectioned, processed for H&E staining and examined by a pathologist. Treatment conditions (+: present, -: not present) are indicated in table including the mice which received bacteria and/or Mannose/IPTG. Histopathology and observations are described in the table. Giemsa staining was performed in an attempt to stain bacteria; none were found in any group (-).

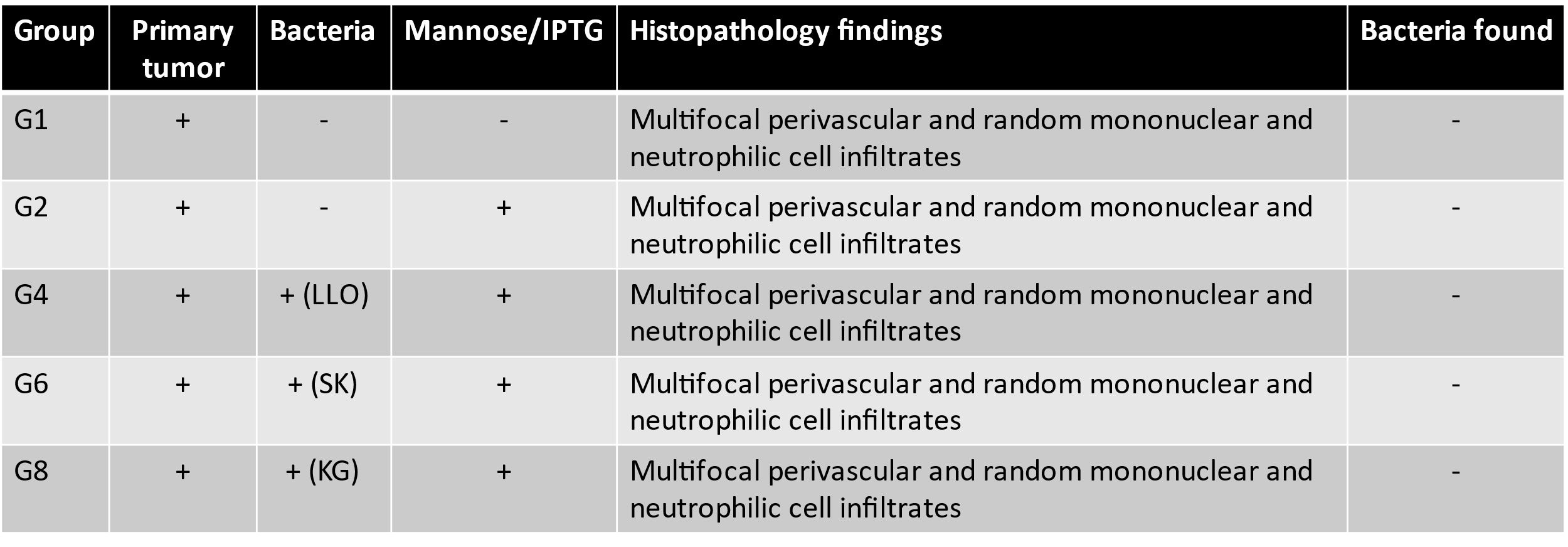
